# Overcoming data limitations in macrogenetics: assessing and accounting for spatial heterogeneity in sequence data

**DOI:** 10.64898/2026.09.29.755480

**Authors:** Maria João Paúl, Matthew O. Moreira, André Vicente Liz, Jeffrey O. Hanson, Pedro Tarroso, Angelica Crottini, Duarte V. Gonçalves, Ana C. Carnaval, Bryan C. Carstens, Craig Moritz, Sílvia B. Carvalho

## Abstract

At large scales, our ability to compare genetic diversity across species is ofien challenged by the difficulty of extracting range-wide sequence data from online databases. Using mitochondrial DNA from amphibians and reptiles spanning across six regions: Brazilian Atlantic Forest, Iberian Peninsula, northwestern Africa, Arabian Peninsula, Madagascar and Australian Monsoonal Tropics, we develop and implement a framework to overcome common limitations in macrogenetics. We compiled and manually curated a spatially explicit genetic dataset from public repositories, quantified the spatial representativeness of genetic sampling across species’ ranges and developed a novel interpolation framework to estimate and map species-specific patterns of intraspecific genetic diversity (ISD) while accounting for heterogeneous spatial sampling. We estimated mean nucleotide diversity across species to produce regional maps for each genetic marker. Our final dataset comprised 182 amphibian and 239 reptile species, encompassing 184 genera from 54 families. Range representativeness of genetic sampling was generally high, despite some variation across study regions. Spatial patterns of ISD revealed geographic heterogeneity, and were consistent across the best-sampled mitochondrial markers. By combining automated and manual georeferencing, quantifying range representativeness, and accounting for bias in spatial interpolations, our pipeline is marker-agnostic and directly applicable to nuclear and genomic data as they become available.

## 1. Introduction

Intraspecific genetic diversity (ISD) is a pillar of biodiversity that underpins the long-term persistence and evolutionary potential of species because it is a prerequisite for adaptation to changing environments (Hoban et al., 2021; Kardos et al., 2021; Sgrò et al., 2011). ISD also supports ecosystem diversity, resilience, function and community structure, thus contributing to ecosystem services pivotal to human well-being (Heuertz et al., 2023; Hoban et al., 2021; Hughes et al., 2008; Sgrò et al., 2011). Human-driven habitat degradation and climatic change are accelerating the erosion of ISD (Leigh et al., 2019; Shaw et al., 2025) through population range shifts and bottlenecks. Understanding where genetic diversity is located across species’ ranges is therefore essential for identifying areas of evolutionary potential, detecting spatial gaps in biodiversity knowledge and informing conservation planning.

The availability of genetic data required to estimate ISD has increased significantly in the past few decades (Blanchet et al., 2017). This is mainly due to the implementation of mandatory data archiving policies (Strasser, 2008), which have led to data deposition in publicly accessible repositories, such as the United States National Institutes of Health genetic sequence database (NCBI, https://www.ncbi.nlm.nih.gov/genbank/), the European Nucleotide Archive (https://www.ebi.ac.uk/ena/browser/home) and the Barcode of Life Data Systems (BOLD, https://boldsystems.org/). This broader availability has been further enhanced by the development of new tools that facilitate repository surveying and data curation and assembly. For instance, Miraldo et al. (2016) developed a pipeline to compile data from NCBI and BOLD that incorporates automated spatial referencing, a procedure similar to that implemented in the phylogatR database (Pelletier et al., 2022; also see Román-Palacios, 2023). Other applications, like SuperCRUNCH (Portik & Wiens, 2020), support automated alignment of large sequence datasets. Together, increasing data availability and computational advances allow researchers to repurpose data within novel macrogenetic frameworks, including inferring phylogeographic patterns at broad taxonomic and spatial scales (Leigh et al., 2021), testing hypotheses about the drivers of genetic variation (Blanchet et al., 2017; Leigh et al., 2021) and informing spatial conservation strategies by explicitly integrating ISD (Carvalho et al., 2017; Nielsen et al., 2023; Schmidt et al., 2025).

Despite these recent advances, substantial limitations remain (Schmidt et al., 2023). Sequence data are still scarce for most taxa, with only 1 to 5% of named species being associated with any genetic data (Hoban et al., 2024), most of which describes variation in mitochondrial DNA (mtDNA). The global distribution of these data remains uneven, with some regions lacking consistent sampling (Csilléry et al., 2025; Miraldo et al., 2016; Paz-Vinas et al., 2025). Geographic information linked to sequence data is also limited: as of 2016, over 85% of all NCBI sequences were missing coordinates (Miraldo et al., 2016). In some cases, such data may be available elsewhere (*e.g.*, Pelletier et al., 2022), but many records exist only in inconsistent and difficult-to-access formats (*e.g.*, supplementary material of scientific publications or tables within the main text). Moreover, as different genetic markers can yield distinct perspectives on the spatial distribution of ISD and its underlying drivers, marker choice can affect comparative analyses (Larkin et al., 2023). Therefore, mtDNA, while having well-documented limitations as a proxy for genome-wide diversity (Galtier et al., 2009), remains by far the most abundant and geographically referenced sequence data available for most taxa. Population-level nuclear, microsatellite and especially genomic data, while increasingly generated, are still too heterogeneously deposited across repositories and too sparse taxonomically and geographically for the kind of multi-region, multi-taxon spatial analyses developed here (Csilléry et al., 2025; Schmidt et al., 2023). As such, mtDNA provides a natural and pragmatic test case for developing macrogenetic pipelines. Ultimately, it remains unclear to what extent the existing data effectively represent genetic diversity across species’ ranges (Miraldo et al., 2016). Addressing these taxonomic, spatial, and marker-specific gaps is crucial to maximize the accuracy of ISD inferences.

Methodologies for mapping broad-scale patterns in ISD must overcome limitations in the underlying sequence data (Leigh et al., 2021). A first assessment by Miraldo et al. (2016) subdivided the globe using a large spatial grid, calculated mtDNA nucleotide diversity for each species within each grid cell and then calculated the average nucleotide diversity in each grid cell. Although this approach successfully accommodated multiple species and provided a framework for subsequent research, uneven sampling across species’ ranges may bias spatial patterns and, in turn, skew results toward locations with greater taxonomic representation and sampling effort. Given that different genetic markers are often used across species, diversity maps also tend to be based on a single marker (usually the one with greater sampling) and are, thus, taxonomically and spatially incomplete, making it difficult to compare spatial patterns observed across studies. Although geostatistical interpolation and modelling techniques such as inverse distance weighting, thin-plate splines or kriging could overcome these limitations (Bishop et al., 2023; Miller, 2005; Vandergast et al., 2011), they have largely been overlooked for mapping broadscale patterns in ISD, mostly due to scarcity of georeferenced data. Consequently, four interrelated challenges continue to limit spatial macrogenetic inference: (i) incomplete georeferencing, (ii) unknown range-wide sampling representativeness, (iii) spatially biased sampling effort, and (iv) the difficulty of converting sparse genetic observations into continuous estimates of diversity across species’ ranges.

Here, we develop and implement a new methodological approach to address challenges in mapping ISD and reduce the existing knowledge shortfall in amphibians and reptiles. We apply this methodological approach across the Brazilian Atlantic Forest (hereafter Brazilian AF), the Iberian Peninsula, northwestern Africa (hereafter NW Africa), the Arabian Peninsula, Madagascar and the Australian Monsoonal Tropics (hereafter AMT). These regions encompass contrasting biogeographic, climatic and evolutionary contexts while spanning broad gradients in species richness, endemism, climatic stability and environmental heterogeneity. Importantly, they also contain some of the most comprehensive and geographically representative genetic datasets currently available for amphibians and reptiles, supported by well-resolved phylogeographic studies across multiple co-occurring taxa. Together, these characteristics provide an ideal framework for developing and evaluating methodological approaches for large-scale macrogenetic analyses. In addition, the long-term research programmes conducted by our team across these regions have provided extensive expertise on their taxonomy, biogeography and evolutionary history, enabling rigorous data curation and interpretation.

To analyse these data, we expanded the approach developed by Miraldo et al. (2016) through three main innovations. First, we implemented a dual retrieval strategy, combining automated data compilation with manual search of georeferenced data in the literature to enhance the completeness of the dataset. Second, we developed a method to quantify the species range representativeness of a set of genetic samples. Third, we introduced a novel interpolation protocol designed explicitly to account for sampling heterogeneity when estimating nucleotide diversity across species ranges.

Using these approaches, we generated maps of ISD for each region. Nuclear data available for the study taxa was sparse, thus, inferred maps only include mtDNA. These insights allow us to discuss the major challenges and opportunities associated with genetic data repurposing for macrogenetic and conservation planning. More broadly, this framework provides a reproducible and transferable approach for improving spatial genetic diversity assessments from heterogeneous public datasets and can be readily extended to nuclear and genomic markers as these become more consistently available.

## 2. Methods

### 2.1. Methodological workflow

To overcome the principal limitations associated with large-scale macrogenetic analyses, we developed a sequential framework that integrates data compilation, quality control and spatial modelling (Figure 1). The workflow begins with the automated retrieval and manual curation of geographically referenced genetic data, followed by the quantification of sampling representativeness across species’ geographic ranges. Genetic sequences are then aligned to estimate species-specific nucleotide diversity, which is subsequently interpolated to generate continuous surfaces of species-level intraspecific genetic diversity. Finally, species-level interpolations are integrated to produce regional ISD maps. Together, these methodological components provide a reproducible framework for transforming heterogeneous public genetic datasets into spatially explicit estimates of intraspecific genetic diversity while addressing incomplete georeferencing, uneven sampling effort and variable sampling coverage across species’ ranges.

**Figure 1.**
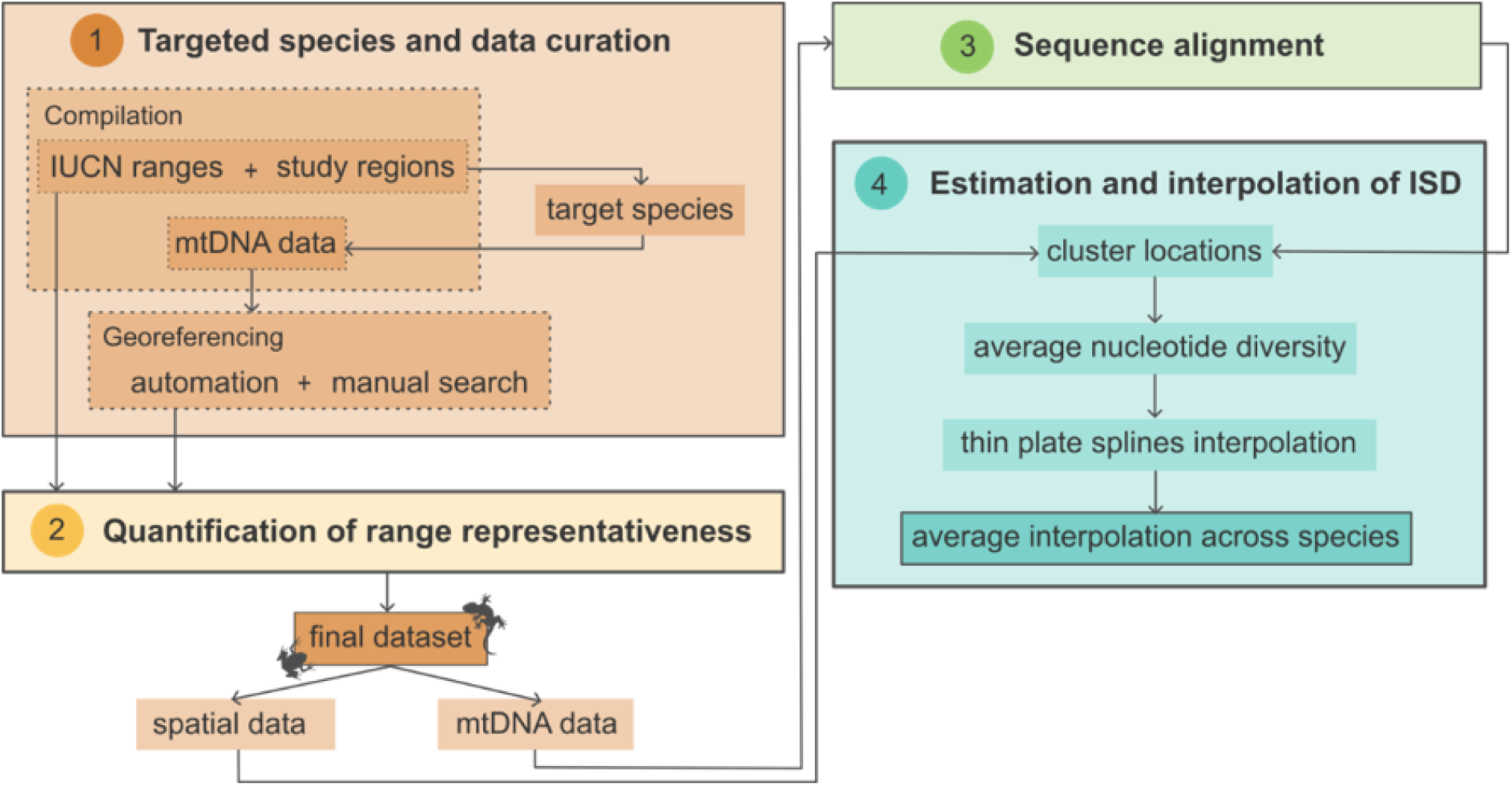
Workflow outlining the main steps of the study pipeline, including data compilation and georeferencing, quantification of range representativeness, sequence alignment, and estimation and interpolation of spatially explicit ISD.

### 2.2. Targeted species

To develop and evaluate our framework across a broad range of evolutionary and biogeographic contexts, we selected all native terrestrial reptile and amphibian species listed in the IUCN Red List of Threatened Species (IUCN, 2022) whose geographic ranges (obtained from the IUCN) intersected any of our six study regions, regardless of threat status. This inclusive approach ensured that the framework was evaluated across a broad taxonomic coverage, encompassing species spanning wide gradients in geographic range size, sampling completeness and phylogeographic complexity.

### 2.3. Genetic data compilation and curation

To maximise the completeness of the genetic dataset, we combined automated retrieval of publicly available sequence data with targeted manual curation. First, we retrieved all mtDNA data available from the NCBI repository (https://www.ncbi.nlm.nih.gov/) through an application-platform interface (API; Entrez Programming Utilities for NCBI), using customized scripts and the ‘rentrez’ package (Winter, 2017) in the R programming environment (R Core Team, 2024). Data were retrieved in July 2022 and February 2023 for amphibians and reptiles, respectively. Since the taxonomy in NCBI can often be misidentified or outdated, we used the Integrated Taxonomic Information System (ITIS) to search for synonyms of each species, using the R package ‘taxize’ (Chamberlain & Szöcs, 2013). For each species and its synonyms, we retrieved all available sequences in FASTA format and associated metadata, which were then organized into data frames using custom R scripts. NCBI metadata retrieved for each sequence included the accession number, species name, entry title, fragment annotation, sequence length, respective isolate or voucher information, and bibliographic information—such as authors, title, journal, and year. When available, we also retrieved associated spatial information, either as geographic coordinates, geographic descriptors (country, locality), or both.

To ensure taxonomic and marker consistency across the compiled dataset, we created a list of alternative names for commonly sequenced markers and standardized their names accordingly. Using taxonomic synonyms, we removed taxonomic duplicates and harmonized all records under a consistent species name. To this end, we identified identical sequences across species and reassigned them based on whether their collection locations fell within the known geographic range of each species or, when necessary, based on expert judgment within our team. We then quantified the total number of sequences per genetic marker for each species. Species were retained for further analysis if they presented at least 10 sequences for at least one mitochondrial marker.

Because a large proportion of publicly available sequences lack complete geographic metadata, automated repository searches alone often fail to recover sufficient information for spatial analyses. Therefore, for the retained species, we complemented NCBI geographic metadata with a thorough manual search of location records. First, we used the previously retrieved bibliographic metadata to perform an automated search for the corresponding Digital Object Identifiers (DOIs) associated with each sequence. We then reviewed each publication and its supplementary materials to extract any additional geographic data related to the sequences (*e.g.*, locality names or coordinates). We verified whether the records referred to individual specimens or haplotypes. In cases where the same haplotype was reported from multiple locations, we duplicated the haplotype entry in our database to reflect its presence in each reported locality. For all sequences with available locality descriptors, we converted these into geographic coordinates using the Google Maps API, accessed through the ‘ggmap’ R package (Kahle & Wickham, 2013). Each locality search was filtered by country to avoid retrieving similarly named localities from different countries. Coordinates assigned to each sequence were prioritized by level of certainty: first, those already available in NCBI; second, coordinates explicitly reported in publications; and third, those inferred from locality descriptors. To further ensure accuracy of the geographic data, we assessed the spatial concordance between the sequence coordinates and the corresponding IUCN species range maps.

Recognizing inherent uncertainty in IUCN range delineations (Liz et al., 2025), we applied species-specific buffer zones to each IUCN range. Given that larger range sizes generally tend to exhibit greater spatial uncertainty, buffer sizes were scaled according to the area of each species’ range (in km²), allowing larger buffers for larger species ranges. Specifically, we defined ten buffer classes according to species range size (km²): 0–100 (10 km), 101–400 (20 km), 401–900 (30 km), 901–1,600 (40 km), 1,601–2,500 (50 km), 2,501–3,600 (60 km), 3,601–4,900 (70 km), 4,901–6,400 (80 km), 6,401–8,100 (90 km), and >8,100 (100 km). Sequences with coordinates falling outside these buffered ranges were excluded. To further refine the dataset, we used the ‘CoordinateCleaner’ R package (Zizka et al., 2019) to remove potentially erroneous coordinates, specifically those located at country centroids, with identical absolute latitude and longitude, near biodiversity institutions, or in marine areas.

### 2.4. Quantification of range representativeness

Because publicly available genetic datasets are often spatially clustered and rarely provide uniform coverage across species’ distributions, we developed a metric to quantify the geographic representativeness of the available sampling. This metric estimates the extent to which sampled localities capture the spatial distribution of a species and provides an objective measure of the confidence that can be placed on subsequent estimates of nucleotide diversity. To assess the sampling representativeness, we used a modified method of the Dynamic Match Coefficient (Sporbert et al., 2019). For each species, species’ ranges were rasterized at 20 progressively coarser grid sizes. Because species differed markedly in geographic range size, raster resolution was scaled proportionally to the extent of each species’ distribution. The maximum pixel size was defined as the square root of the species’ range area multiplied by the initial pixel side length (*i.e.*, 0.008333°, ∼1km), while the minimum pixel size was set to one-twentieth of that value. At each spatial resolution, we calculated (i) observed representativeness, (ii) expected representativeness, and (iii) relative representativeness.

To quantify observed representativeness (i), we employed the urap.proportion.held function from the ‘raptr’ R package (Hanson et al., 2018), which computes the proportion of total variance explained by the observed sampling points relative to a uniform distribution across the species’ range. To calculate expected representativeness (ii), we employed the same function but replaced the observed sampling points with an evenly distributed set of points across the species’ range. This was achieved by performing k-means clustering on a set of 10,000 random points to generate a number of clusters equal to the number of observed sampling points. The centroids of these clusters, calculated using Voronoi polygons in the ‘dismo’ R package (Hijmans et al., 2024), were treated as evenly distributed sampling points. To compute relative representativeness (iii), we calculated the ratio of observed (i) to expected (ii) representativeness. Finally, we calculated the area under the curve (AUC) for the relative representativeness (iii) values across all pixel sizes (Figure S1). This index was normalized to range from zero to one, and the resulting metric was used to quantify range representativeness in subsequent analyses. Values approaching one indicate that sampling is broadly distributed across the species’ geographic range, whereas values approaching zero indicate spatially clustered or geographically unrepresentative sampling. By summarising relative representativeness as the area under the curve across multiple raster resolutions, the index provides a scale-independent measure that is robust to the arbitrary choice of pixel size.

### 2.5. Sequence alignment

To ensure consistent and reproducible sequence alignments across the large number of species and genetic markers analysed, we implemented a semi-automated alignment workflow using the software SuperCRUNCH, a python-based toolkit specifically developed for assembling and aligning large phylogenetic datasets (Portik & Wiens, 2020). The workflow first verified taxonomic identities and automatically separated sequences into marker-specific FASTA files whenever multiple loci were present.

In the ‘Sequence Filtering’ step, sequences were clustered by similarity and homologous regions were identified. Because each dataset contained sequences from a single species, we used the module ‘Cluster_Blast_extract.py’ to refine homologous regions. We standardized the direction of all sequences in each marker-specific FASTA file and aligned the sequences using the MAFFT algorithm. All alignments were subsequently inspected manually to ensure accuracy. Sequences that could not be reliably aligned were excluded from subsequent analyses.

### 2.6. Estimation and interpolation of ISD

For each species and mitochondrial marker, we estimated nucleotide diversity across the overall species’ geographic range. Species were excluded from the analysis if sequence data were available from fewer than five distinct locations or if their relative representativeness was zero, indicating extremely spatially clustered sampling.

Because genetic sampling is often highly clustered geographically, direct interpolation from all sampling locations would disproportionately weight densely sampled areas and potentially bias estimates of nucleotide diversity. To reduce this effect while retaining all available genetic information, we developed a spatial subsampling procedure based on clusters and bins. Sampling locations were first grouped into spatial clusters by applying hierarchical clustering (WPGMA) to a pairwise Euclidean distance matrix calculated from the geographic coordinates of all sampling locations. The clustering threshold was defined as the first quartile of the pairwise distance distribution to balance two competing objectives: reducing spatial redundancy by grouping nearby sampling locations likely to contain redundant spatial information while preserving sufficient geographic subdivision across the species’ range. The resulting clusters were then used to construct a series of spatially balanced sampling datasets (hereafter referred to as bins). The largest cluster determined the number of bins to be generated. Each sampling location within this cluster was assigned to a different bin, whereas one sampling location from every remaining cluster was allocated to each bin. For clusters containing fewer sampling locations than the total number of bins, individual locations were assigned to multiple bins, ensuring that every bin contained one representative from every cluster while every sampling location contributed to at least one bin. Consequently, each bin represented an alternative, geographically balanced realization of the available sampling, reducing the influence of locally intensive sampling without discarding genetic information.

For each sampling location within each bin, we calculated nucleotide diversity using the ‘pegas’ R package (Paradis, 2010). We defined a local neighbourhood around each sampling location with a radius equal to the maximum nearest-neighbour distance, thereby ensuring that each estimate was based on at least two sequences. Within each neighborhood, we identified haplotypes and computed all pairwise haplotype combinations. Nucleotide diversity was calculated as the mean across all pairwise comparisons rather than from the pooled neighbourhood dataset. This approach reduced biases associated with heterogeneous local sampling effort while producing geographically comparable estimates of nucleotide diversity. To estimate continuous spatial patterns of nucleotide diversity across each species’ geographic range, we fitted a thin plate splines model using the ‘fields’ R package (Nychka et al., 2021) to the nucleotide diversity estimates at each sampling location and calculated the associated standard errors. Thin plate splines were selected because they do not require the estimation of species-specific variograms, which would be difficult to estimate robustly across hundreds of species with heterogeneous sample sizes and irregular spatial sampling. This characteristic makes thin plate spines particularly suitable for analysis of large comparative macrogenetic datasets, while also providing standard error estimates that quantify interpolation uncertainty. The interpolation was performed independently for each bin, generating multiple spatial predictions based on alternative geographically balanced representations of the available sampling. Final nucleotide diversity and associated standard error surfaces were obtained by averaging predictions across all bins, thereby reducing the influence of uneven sampling while preserving broad-scale spatial patterns of genetic diversity. Because mitochondrial markers differ substantially in their evolutionary rates and absolute levels of nucleotide diversity, direct comparisons among markers are inappropriate. We therefore applied min-max normalization to the nucleotide diversity values, rescaling each marker independently to a common range between zero and one while preserving its relative spatial variation. We then cropped each species’ interpolated nucleotide diversity surface to the boundaries of the region intersected by its range (Brazilian AF, Iberian Peninsula, NW Africa, Arabian Peninsula, Madagascar and AMT). Finally, regional maps of intraspecific genetic diversity were generated by calculating, for each mitochondrial marker, the mean normalized nucleotide diversity across all species occurring within each study region.

## 3. Results

### 3.1. Number of species per region

We found 182 amphibian species (17.5% of the 1040 initial target amphibians; Dataset S1) and 239 reptile species (18.8% of the 1268 initial target reptiles; Dataset S1) presenting sufficient data available to map patterns of ISD (*i.e.*, 421 species in total). As such, of the 2308 initial target species occurring among our study regions—with some species occurring in multiple regions—only 421 species (Brazilian AF: 102/771; Iberian Peninsula: 56/78; NW Africa: 70/173; Arabian Peninsula: 63/199; Madagascar: 85/695; AMT: 72/543) met our inclusion criteria: at least 10 sequences for at least one mtDNA marker, samples from at least five distinct locations, and a representativeness value greater than zero. These 421 species formed the dataset used in all subsequent analyses.

These taxa encompassed 184 genera from 54 families, representing 38% and 72% of the 489 genera and 75 families identified across the 2308 initial target species, respectively. We also found that regions that had a greater number of species tended to have greater data scarcity. For example, the Brazilian AF, Madagascar and AMT tended to have the smallest proportion of amphibian and reptile species with sufficient data available for mapping ISD (Brazilian AF: 13%, Madagascar: 12%, AMT: 13%). Alternatively, the Iberian Peninsula consistently had the largest proportion (72%) of species with sufficient data available for mapping ISD. Our findings demonstrate that despite substantial improvements in data compilation and georeferencing, considerable variation in both taxonomic representation and geographic sampling coverage remained across the study regions.

### 3.2. Number of sequences per genetic marker

Overall, a total of 73,256 mitochondrial sequences corresponding to our final taxa were assembled. Only 11% (n=8,121) of these sequences were originally georeferenced in NCBI. Through manual compilation of geographic coordinates and/or locality data from publications, and by retrieving geographic data from locality descriptors of associated metadata, we increased the total proportion of georeferenced data to 91% (n=66,654). The number of georeferenced sequences varied substantially among genetic markers in both taxa. For amphibians, 16S and CYTB were the most available markers, and for reptiles, CYTB and ND4 predominated (Figure 2).

**Figure 2.**
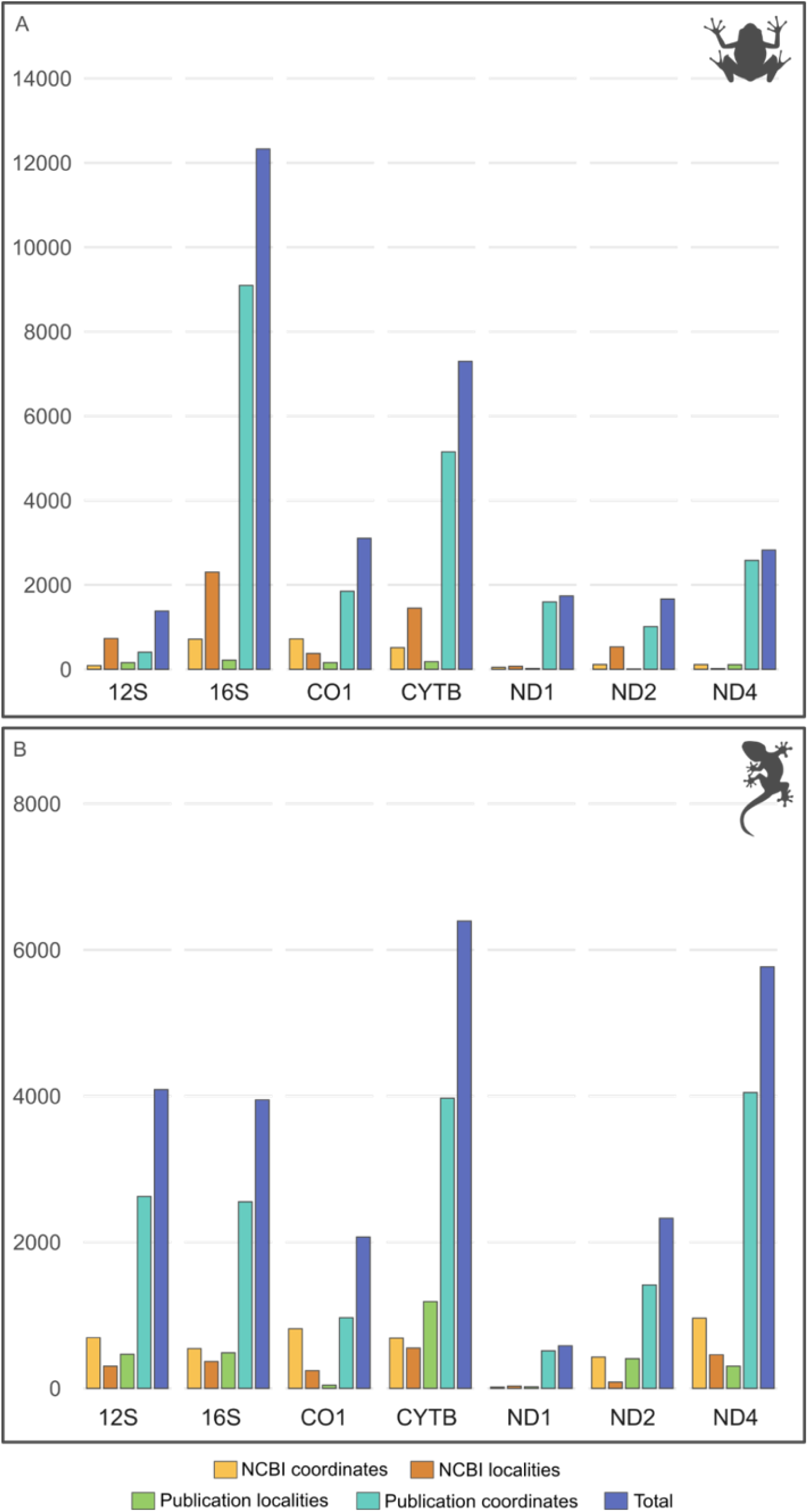
Barplots of number of georeferenced sequences for each genetic marker according to the source of geographic data for amphibians (A) and reptiles (B) after applying the coordinate priority framework (for more details, see Methods section). Colored bars indicate the different sources of occurrence data: NCBI coordinates, NCBI localities, publication localities, publication coordinates, and the overall total. Bars represent the final georeferenced data used for downstream analyses, obtained after conducting spatial concordance analysis.

### 3.3. Range representativeness

Species with high range representativeness show sampling localities distributed broadly across their geographic range, regardless of species’ range size (Figure S2). In contrast, species with low range representativeness showed sampling locations in a small portion of their distribution range with spatial clustering (Figure S3). This highlights how similar range representativeness values can occur in species with very different range sizes.

Range representativeness varied similarly across study regions in both amphibians and reptiles (Figure 3). Range representativeness was generally highest for species occurring in the Iberian Peninsula and NW Africa. Regions with a greater number of species, such as the Brazilian AF, Madagascar and AMT, tended to have greater representativeness variability. The Arabian Peninsula displayed more contrasting patterns between amphibians and reptiles, with substantially higher number of species and variability in reptiles than amphibians.

**Figure 3.**
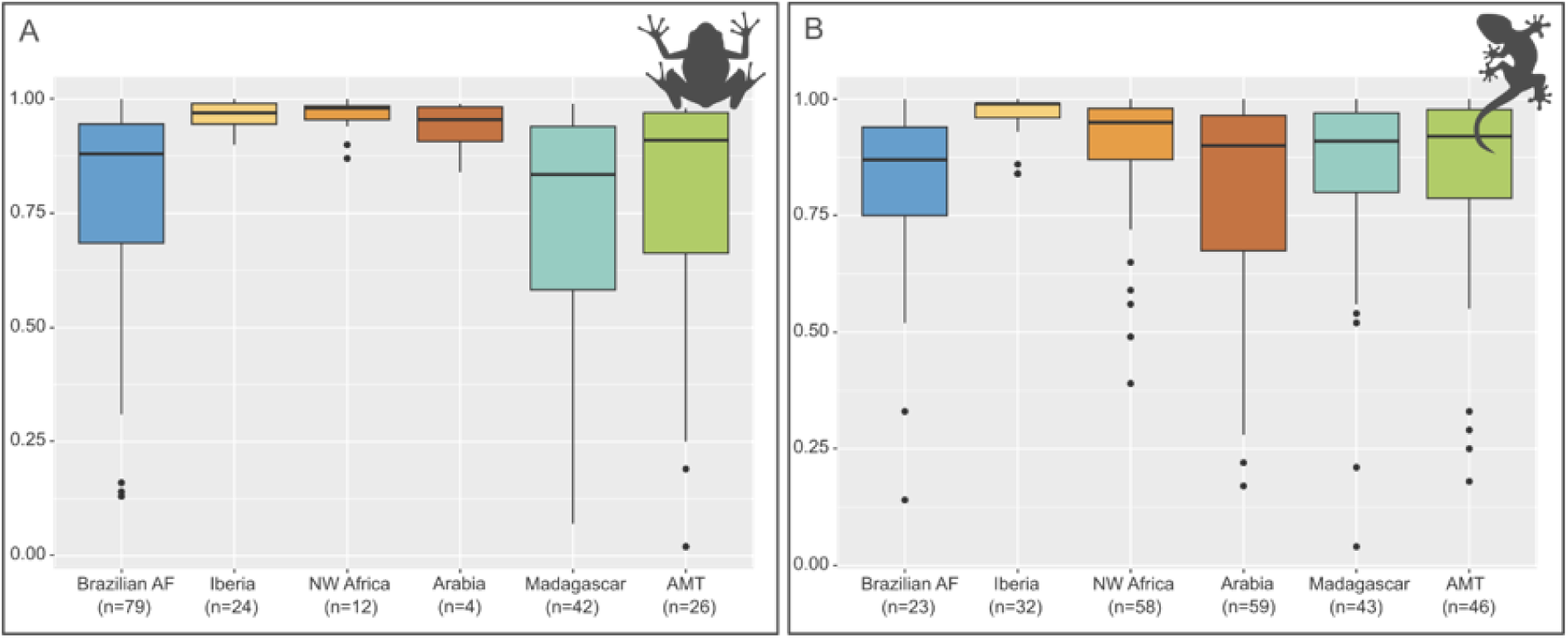
Boxplots of range representativeness per study region in amphibians (A) and reptiles (B). Each box–whisker plot indicates the median (bold lines) and the interquartile range (boxes). Colours represent regions of occurrence.

Range representativeness was positively correlated with the number of georeferenced sequences in both amphibians and reptiles (Figure S4), although considerable variation was observed among species with similar numbers of sequences. Amphibians also exhibited a positive relationship between representativeness and geographic range size, whereas no clear relationship was observed for reptiles (Figure S5).

### 3.4. Spatial patterns of ISD

The proposed workflow generated continuous ISD surfaces for all six study regions despite marked differences in taxonomic richness, sampling density and marker availability. Mean nucleotide diversity estimates based on the mitochondrial marker available for the largest number of species in each region revealed pronounced spatial heterogeneity in both amphibians and reptiles (Figures 4–5; Figures S6–S29 present the results for all markers). Associated standard error surfaces were generated for every interpolation, providing spatially explicit estimates of prediction uncertainty (Figures S6–S29).

**Figure 4.**
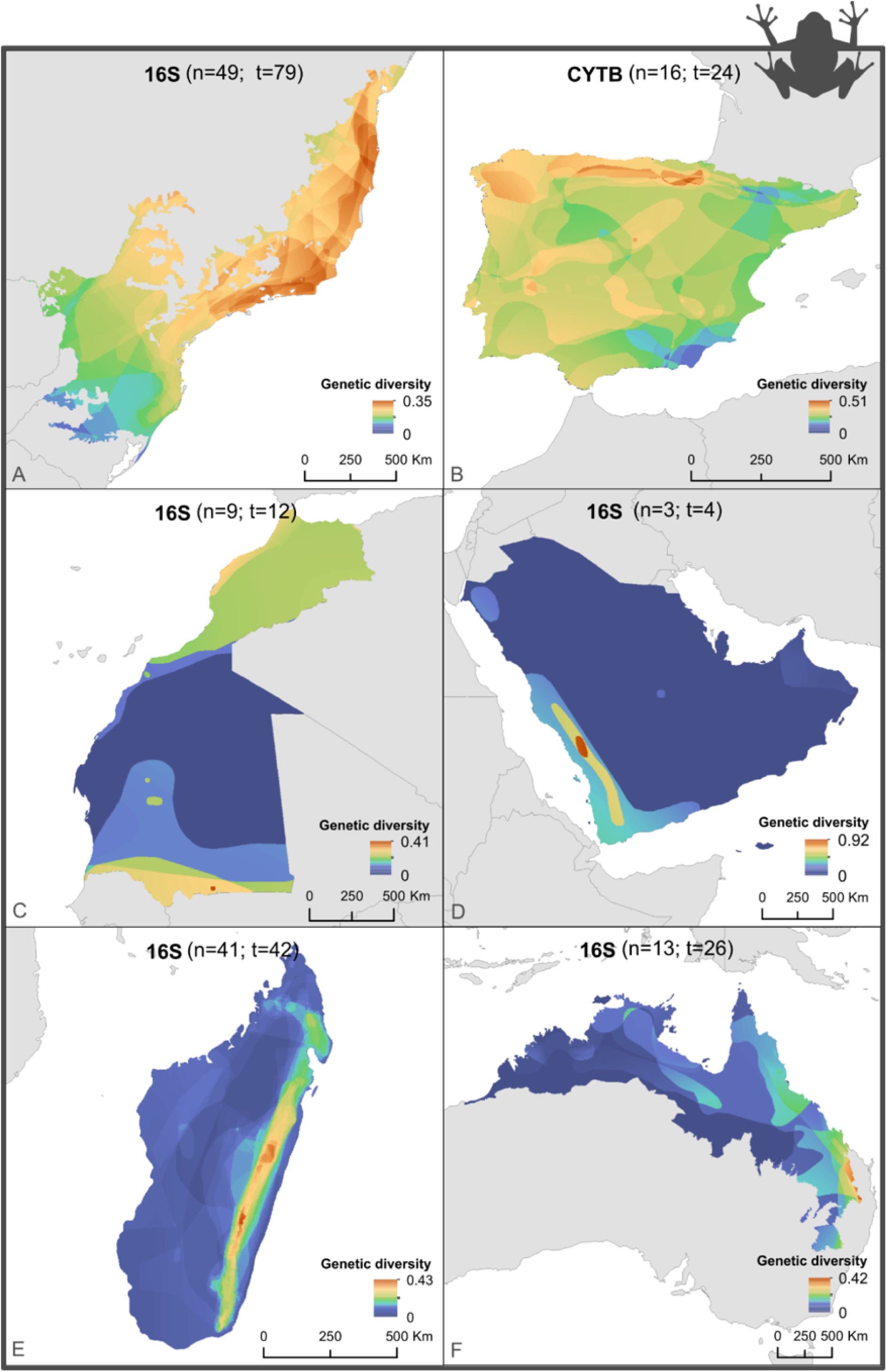
Mean amphibian nucleotide diversity for the genetic marker with the highest number of species in each study region. (n) is the number of species represented by the plotted marker, (t) is the total number of species retained in the final dataset for that region: (A) Brazilian AF, (B) Iberian Peninsula, (C) NW Africa, (D) Arabian Peninsula, (E) Madagascar and (F) AMT.

**Figure 5.**
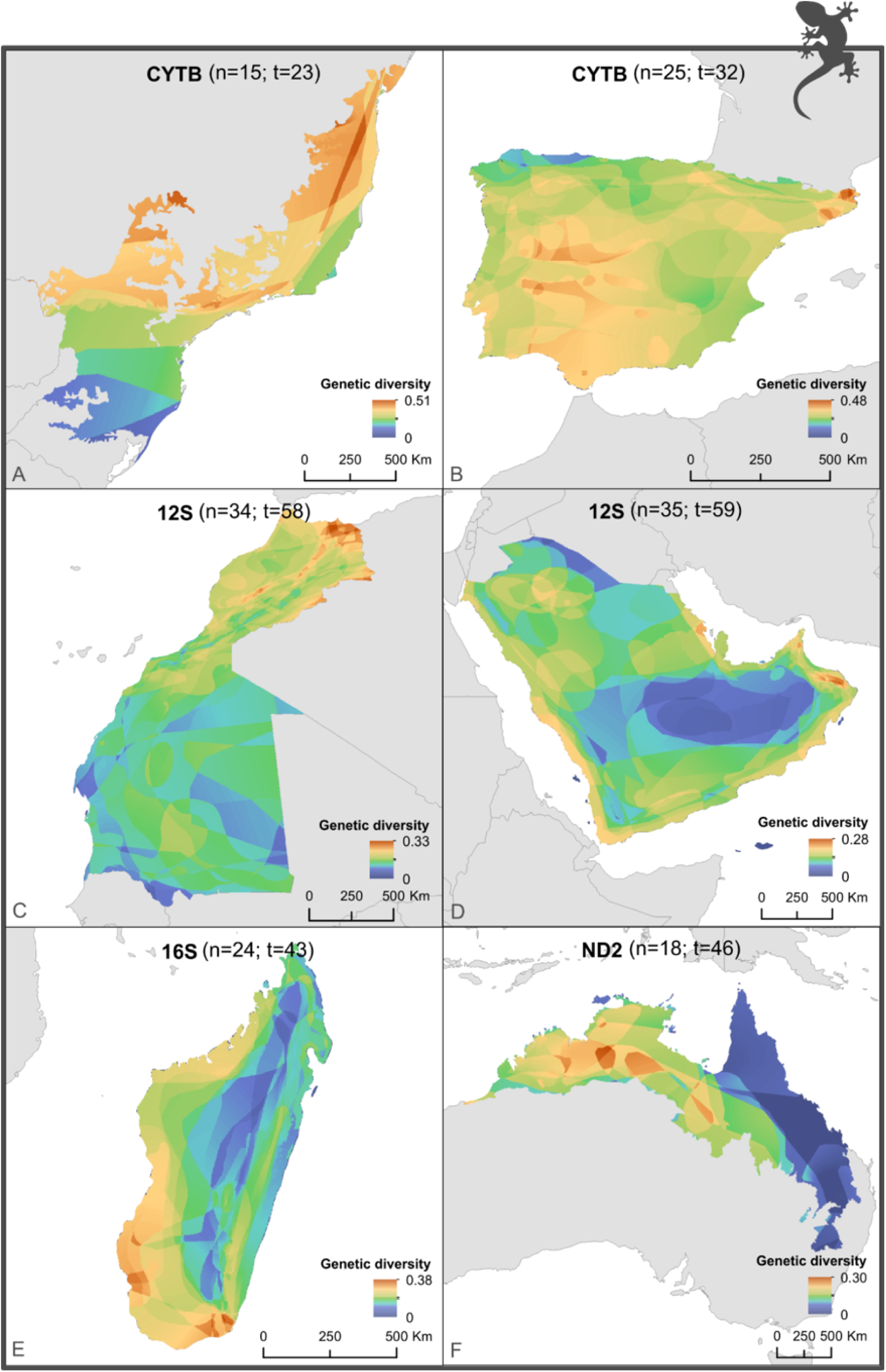
Mean reptile nucleotide diversity for the genetic marker with the highest number of species in each study region. (n) is the number of species represented by the plotted marker, (t) is the total number of species retained in the final dataset for that region: (A) Brazilian AF, (B) Iberian Peninsula, (C) NW Africa, (D) Arabian Peninsula, (E) Madagascar and (F) AMT.

For amphibians (Figure 4), the Brazilian AF exhibited strong spatial structuring, with lower mean ISD in the southernmost portion of the forest. The Iberian Peninsula showed a similarly heterogeneous pattern, with higher mean ISD along its northern range. NW Africa and the Arabian Peninsula, the regions with the least number of amphibian species, showed localized hotspots in the northwestern and southern regions of NW Africa, and in southwestern Arabian Peninsula. Madagascar and AMT exhibited some spatial heterogeneity, with higher ISD occurring in the central-eastern region of Madagascar and along the central-eastern coast of Australia.

For reptiles (Figure 5), the Brazilian AF also exhibited strong spatial structuring, but with higher ISD along the northeastern and central regions. Likewise, the Iberian Peninsula showed spatial heterogeneity, with higher ISD along its central and northwestern ranges. NW Africa and the Arabian Peninsula showed considerable spatial variation of ISD patterns, with higher diversity occurring in the northeast region of NW Africa and along the western coast of the Arabian Peninsula. Madagascar and AMT also exhibited spatial structuring, with higher ISD in the southern and southwestern regions of Madagascar and along the central-northern region of Australia.

## 4. Discussion

This study expands current macrogenetic approaches by presenting an integrated framework for estimating spatial patterns of intraspecific genetic diversity from heterogeneous public genetic datasets. By combining automated and manual geographic data retrieval, a quantitative assessment of sampling representativeness, and an interpolation procedure specifically designed to reduce the effects of uneven geographic sampling, our framework addresses several methodological limitations that have constrained previous large-scale macrogenetic studies. The integration of these complementary components enabled the assembly of a comprehensive spatial genetic dataset comprising more than 66,000 georeferenced mitochondrial sequences from over 400 amphibian and reptile species. The application of this framework across six contrasting biogeographic regions generated continuous spatial estimates of ISD despite substantial differences in taxonomic representation, marker availability and sampling density, illustrating its transferability across diverse ecological and evolutionary settings.

At the same time, our results highlight persistent limitations associated with publicly available genetic datasets and their associated metadata. Despite extensive manual curation and georeferencing efforts, taxonomic representation and geographic sampling coverage remained highly uneven across study regions.

### 4.1. Range representativeness

The range representativeness metric proposed here provides a quantitative assessment of how well available genetic sampling captures the geographic distribution of a species. Although similar concepts have occasionally been explored in biodiversity monitoring, spatial representativeness has rarely been explicitly incorporated into macrogenetic studies, where sampling effort is typically quantified using only the number of sequences or sampling locations (*e.g.*, Amador et al., 2024; Barrow et al., 2021). Such measures provide little information on how well genetic sampling captures the geographic extent of a species’ distribution. An important property of the proposed metric is that it is less sensitive to any single raster resolution and facilitates comparison among species with different range sizes. By explicitly incorporating the spatial distribution of sampling, the proposed metric complements conventional measures of sampling effort and provides a more informative assessment of data quality for large-scale genetic analyses. Beyond its use as a filtering criterion, it could be used to weight species according to sampling reliability in community-level analyses or to identify taxa and regions where additional sampling would most improve estimates of genetic diversity.

Despite differences in taxonomic representation among regions, range representativeness was generally moderate to high for both amphibians and reptiles, indicating that the final datasets captured a substantial proportion of the geographic ranges of the retained species. The positive relationship between representativeness and the number of georeferenced sequences indicates that species with more extensive genetic sampling are generally more likely to achieve broader geographic coverage (Figure S4). However, the considerable variability among species with similar numbers of sequences demonstrates that increasing sequence availability alone does not necessarily improve spatial representativeness. This finding highlights that the number of available sequences should not be interpreted as a surrogate for sampling quality, as extensive genetic datasets may still provide poor representation of a species’ geographic distribution.

The positive relationship between representativeness and geographic range size observed in amphibians (Figure S5) may reflect the greater detectability and broader geographic accessibility of widespread species. In contrast, narrow-ranged species often have smaller population sizes, occur in less accessible areas, or have been more recently described, resulting in less comprehensive genetic sampling. However, this relationship was not evident in reptiles, suggesting that the influence of species’ distributional characteristics on sampling coverage may differ among taxonomic groups.

The greater variability in representativeness observed in the Brazilian AF, Madagascar and the AMT is consistent with well-documented global disparities in genetic data availability, where tropical biodiversity hotspots simultaneously harbour exceptionally high species richness and some of the greatest knowledge shortfalls (Miraldo et al., 2016). Collectively, these findings reinforce the importance of explicitly accounting for spatial sampling coverage when assembling large-scale macrogenetic datasets and identifying priorities for future genetic sampling.

### 4.2. Spatial patterns of ISD

Spatial patterns of ISD revealed regional heterogeneity in amphibians and reptiles. Regions with complex topography and biogeographic histories, such as the Brazilian AF, Madagascar and AMT, harbour deeply structured lineages and high levels of microendemism (Antonelli et al., 2022; Carnaval et al., 2014; Ferreira et al., 2023; Rosauer et al., 2016), which likely contributed to the fine-scale patterns of mtDNA diversity recovered here. Hotspots of ISD across our study regions closely mirror those of species richness (*e.g*., Madagascar, Antonelli et al., 2022) and endemism (*e.g.,* Brazilian AF, Peres et al., 2020), climatic refugia (*e.g.,* Brazilian AF, Carnaval et al., 2009; Iberian Peninsula, Gómez & Lunt, 2007; NW Africa and Arabian Peninsula, Liz et al., 2021; AMT, Schneider & Moritz, 1999), and contact zones or phylogeographic breaks (*e.g.,* NW Africa, Liz et al., 2023; AMT, Fenker et al., 2024). Nevertheless, we note that these estimates can be influenced by differences in taxonomic representation, and geographic sampling effort across regions (Figure S6-S29). Regions with fewer species or limited sampling (Figure S12) may have their diversity patterns underestimated. Uneven sampling within regions can also lead to apparent spatial variation in diversity that reflect sampling effort rather than underlying genetic variation (*e.g.,* Figure S10). Although the representativeness assessment and spatial subsampling procedure were specifically developed to reduce these biases, they cannot fully compensate for major taxonomic and geographic knowledge gaps.

Spatial patterns of ISD were generally consistent across the best sampled mitochondrial markers, though some notable differences were observed (*e.g.,* Figure S8 and S20), underscoring the importance of multi-locus approaches for more robust spatial genetic pattern inference. Because estimates for different markers often included different species and differed in sequence length and sample size, these contrasts may partly reflect sampling and taxonomic composition rather than biological differences among markers. Nevertheless, the broad agreement among the best-sampled markers indicates that the main regional patterns were not driven by a single locus. Still, validation with spatially representative nuclear data remains important.

### 4.3. Mitochondrial DNA in macrogenetics

Mitochondrial ISD patterns can differ from those inferred from genomic data (Schmidt & Garroway, 2021), but mtDNA remains widely used in macrogenetics due to its extensive availability (Leigh et al., 2021). Selection, mutation rate and coalescent variance may influence spatial patterns of mtDNA diversity, potentially distorting interpretations of phylogeographic structure (French et al., 2023). Furthermore, mtDNA cannot provide estimates of key population genetics parameters such as heterozygosity, but the method presented here can easily accommodate nuclear and genomic-level data.

Recent studies and initiatives have tried to overcome limitations of mtDNA use in macrogenetics by aggregating nuclear, microsatellite and genomic data (*e.g.,* Csilléry et al., 2025; De Kort et al., 2021; Yashima & Innan, 2017). However, most nuclear sequence information used for analyses of intraspecific diversity are SNPs whose heterogeneous storing practices across repositories and data types make it difficult to locate, retrieve, and aggregate (Schmidt et al., 2023). Given differences in bioinformatic processing, it is also difficult to compare estimates of diversity across studies. Thus, despite recent efforts, spatially explicit nuclear datasets for a wide variety of taxa remain scarce. As such, large-scale assessments of mtDNA diversity continue to play a key role in advancing our understanding of global spatial patterns of genetic variation (French et al., 2023; Moreira et al., 2026). However, its application and interpretation should be made carefully, particularly at large spatial scales, where biases due to heterogeneous sampling, taxonomic uncertainty and marker-specific constraints are more likely to influence inferred patterns. Multi-marker validation should thus be a priority for future macrogenetic studies as nuclear data becomes more spatially representative.

### 4.4. Data repurposing: major challenges and opportunities

Despite recent advances in repurposing of sequence data, considerable challenges remain. Even though the availability has significantly increased, sequence data remain scarce and uneven across most taxa (Hoban et al., 2024). In addition, many studies either fail to upload their genetic data or only deposit a subset, often limited to unique haplotypes (Paz-Vinas et al., 2021; Pope et al., 2015).

A major limitation identified in this study was the scarcity of geographic metadata associated with publicly available sequence data. As shown in Figure 2, automated retrieval (*e.g.,* NCBI coordinates or NCBI locality metadata) contributed to georeference only a small portion (∼11%) of our data. However, systematic manual curation of coordinates, locality descriptions and associated publications increased this proportion to more than 90%, demonstrating that much of the missing geographic information already exists but remains inaccessible to automated workflows because of inconsistent metadata reporting. This finding corroborates previous findings suggesting that the majority of sequence data in public repositories lack the corresponding geographic data (Miraldo et al., 2016; Pope et al., 2015).

The lack of standardized geographic metadata represents a major obstacle for large-scale macrogenetic studies because it limits the reuse of existing genetic resources and can introduce substantial geographic biases. For example, when coordinates or localities are deposited in a database, it is common for only one of the many locations where a haplotype was found to be recorded (Paz-Vinas et al., 2021). Additionally, geographic data are frequently not linked to NCBI accession numbers which, in turn, introduces challenges to cross-referencing genetic data with genetic data. Furthermore, in cases where geographic data are recorded, they are often stored in different formats (*e.g.*, plain text and various table formats) across different records and databases which, in turn, makes it difficult to automate data extraction. Consequently, integrating genetic and geographic data from multiple studies often requires extensive manual curation and, in some cases, direct communication with authors, making large-scale data compilation extremely time-consuming.

Methodological and taxonomical inconsistencies across studies can also hinder data repurposing efforts. Researchers often use different genetic markers depending on the location and taxa (Figure 2), making it difficult to homogenize and compare the available information. When the same marker is used, there is still substantial variation in the fragment size and marker coverage, due to the use of different primers targeting different portions of the genetic marker across different studies. This leads to difficulties in sequence overlap since there might be different portions of the same genetic marker, making sequence alignment difficult if not impossible. Recent studies have also unveiled hidden errors in voucher data deposited in online repositories (Carné et al., 2025), further complicating the reuse of genetic data. Moreover, differences in sampling effort and coverages can affect spatial pattern detection. Both of these issues, marker inconsistencies and sampling biases, further influence the spatial structure of ISD.

Collectively, these limitations substantially reduce the value of existing genetic resources for macrogenetic analyses. There is an urgent need for a generalized and standardized framework for metadata reporting and repurposing (see Crandall et al., 2023; Forsdick et al., 2025; Leigh et al., 2024). Although the deposition of sequence data and metadata has long been recognized as important, such a framework should include clear guidelines for sequence data and metadata deposition in open-access repositories, including its geographic data. It should also promote the standardization of marker selection and sequence alignment practices. These improvements would certainly help reduce the shortfall of knowledge on spatial ISD patterns and promote its integration in conservation planning.

## Acknowledgements

This work was funded by Fundação para a Ciência e Tecnologia, I.P, via national funds, under the scope of project PLACES - “PLAnning for the Conservation of life on Earth in the light of evolution and Sustainability” (DOI 10.54499/2022.04598.CEECIND/CP1730/CT0006).

MJP was supported by Fundação para a Ciência e Tecnologia through a PhD Grant (SFRH/BD/08664/2020). SBC was supported by Fundação para a Ciência e Tecnologia through an individual scientific employment program contract (CEECIND/CP1730/CT0006). Ana C acknowledges funding by the US National Science Foundation (DBI 1926928, DEB 1343578).

