## Supporting Information for "Overcoming data limitations in macrogenetics: assessing and accounting for spatial heterogeneity in sequence data"

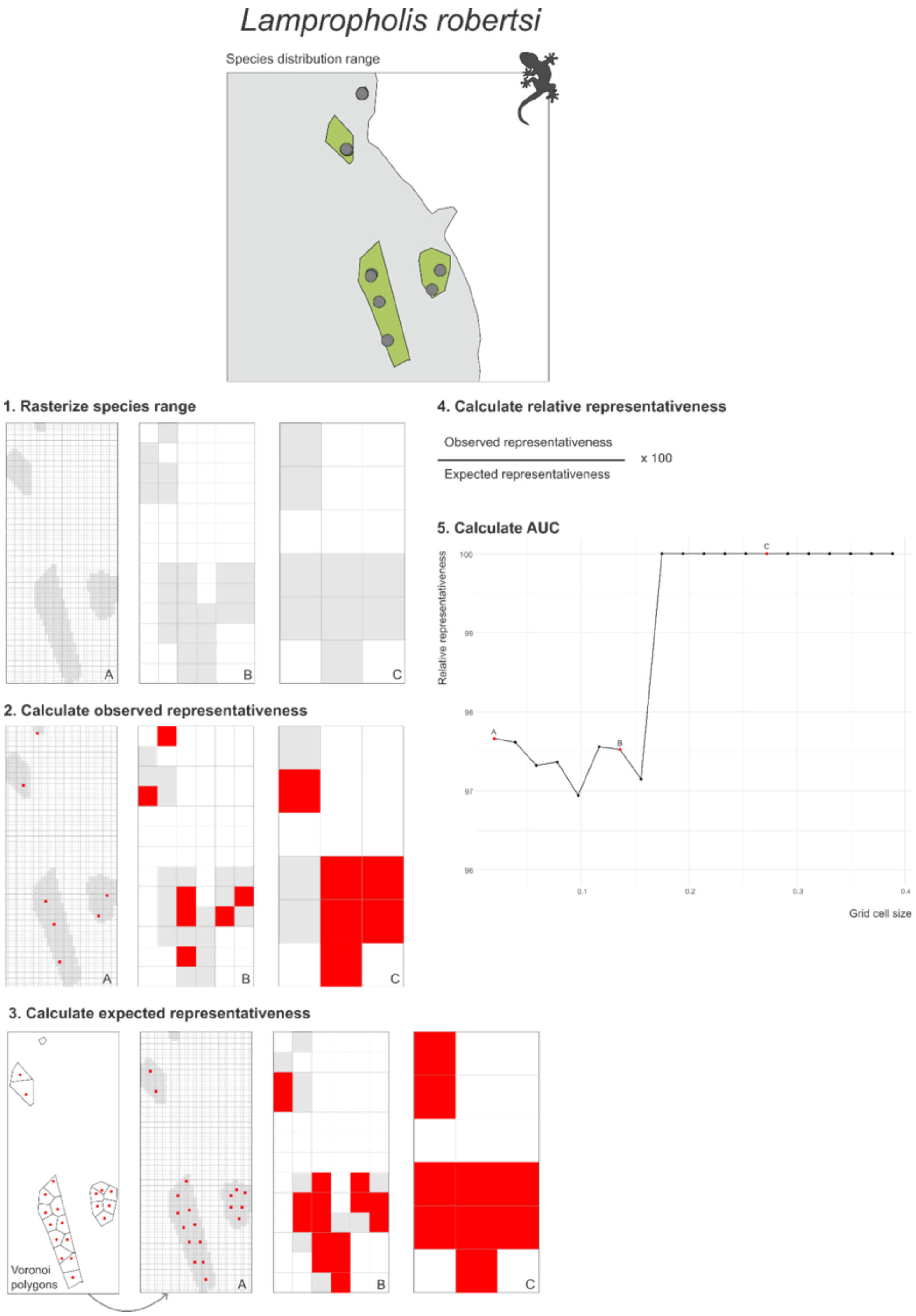


**Figure S1.** Modified DMC method to calculate species' range representativeness, using *Lampropholis robertsi* as example. (A, B and C) illustrate 3 examples—out of the 20 calculated—of different grid cell sizes. Shaded areas represent species' distribution range and red points correspond to georeferenced sampling locations.

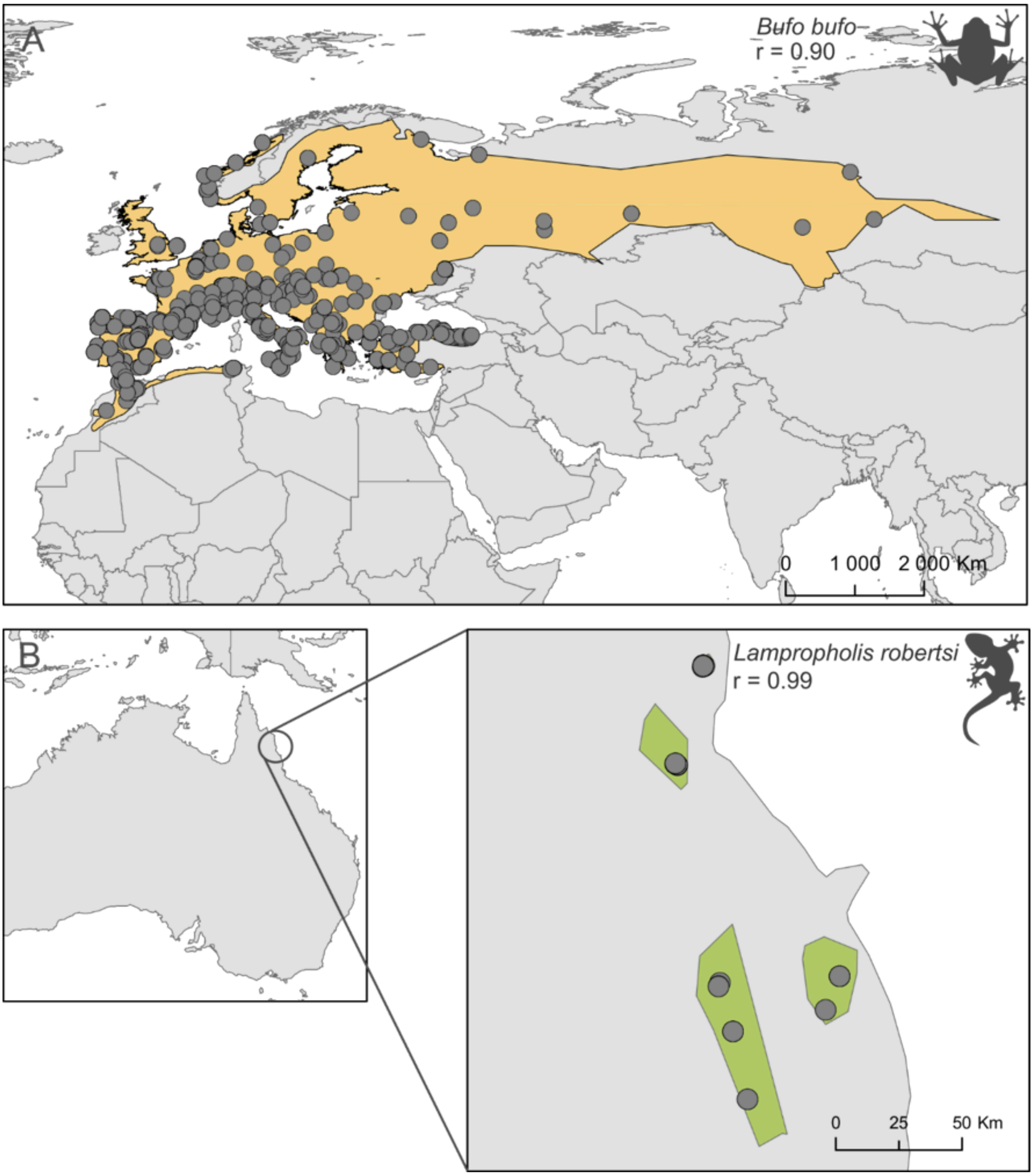


**Figure S2.** Examples of species with high range representativeness (r): (A) wide-ranging amphibian—*Bufo bufo*, and (B) narrow-ranging reptile—*Lampropholis robertsi*. Shaded areas illustrate species' distribution range and points correspond to georeferenced sampling locations.

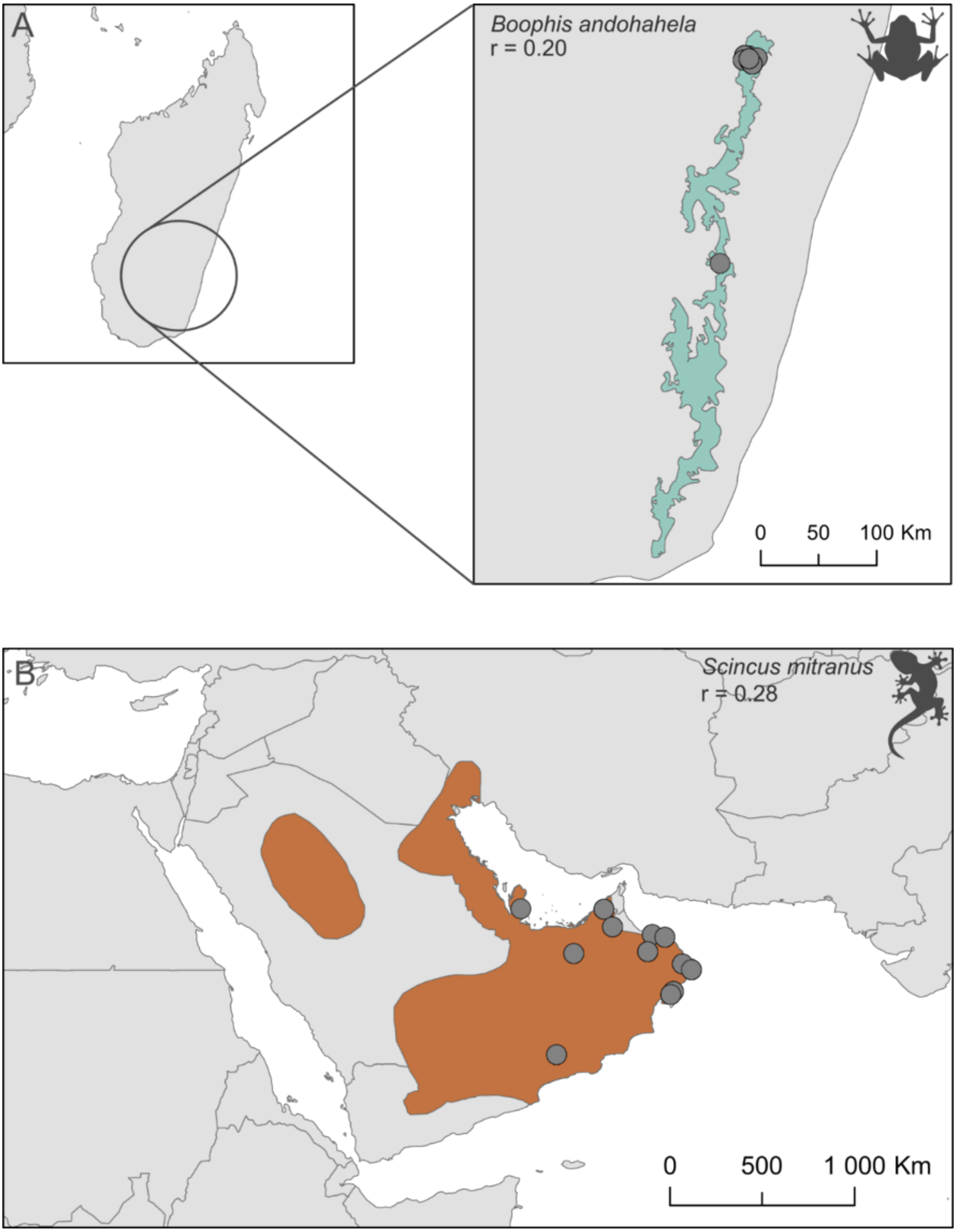


**Figure S3.** Examples of species with low range representativeness (r): (A) narrow-ranging amphibian—*Boophis andohahela*, and (B) wide-ranging reptile—*Scincus mitranusi*. Shaded areas illustrate species′ distribution range and points correspond to georeferenced sampling locations.

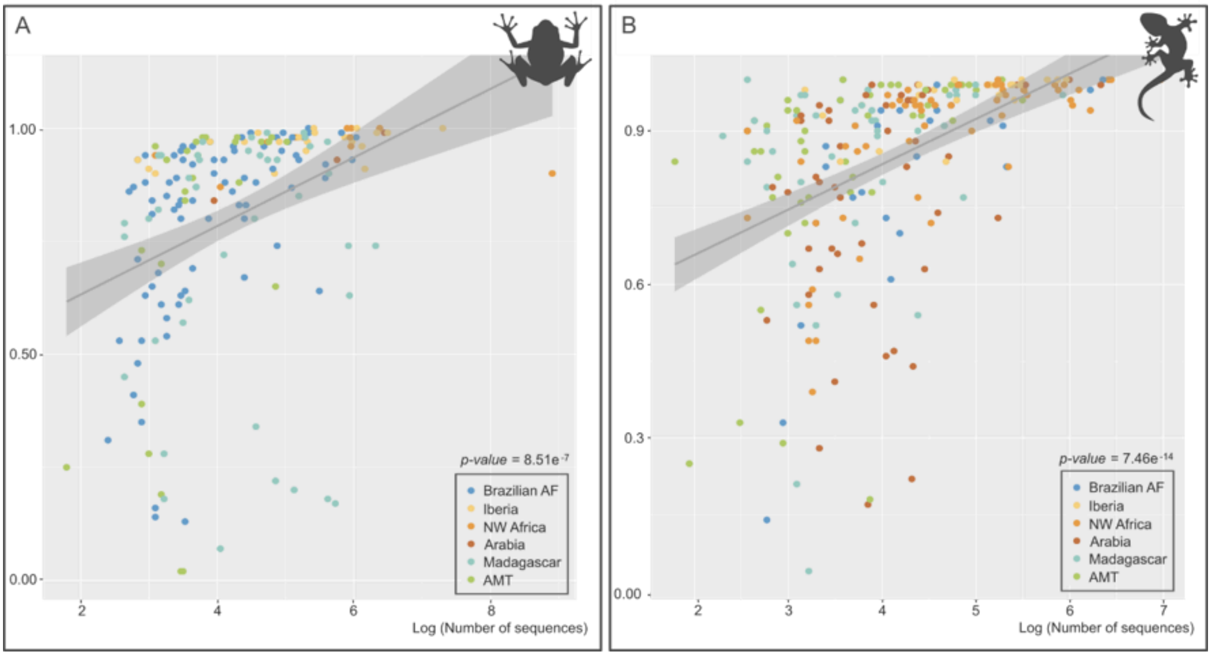


**Figure S4.** Relationship between range representativeness and log-transformed number of georeferenced sequences in amphibians (A) and reptiles (B). Colours represent regions of occurrence.

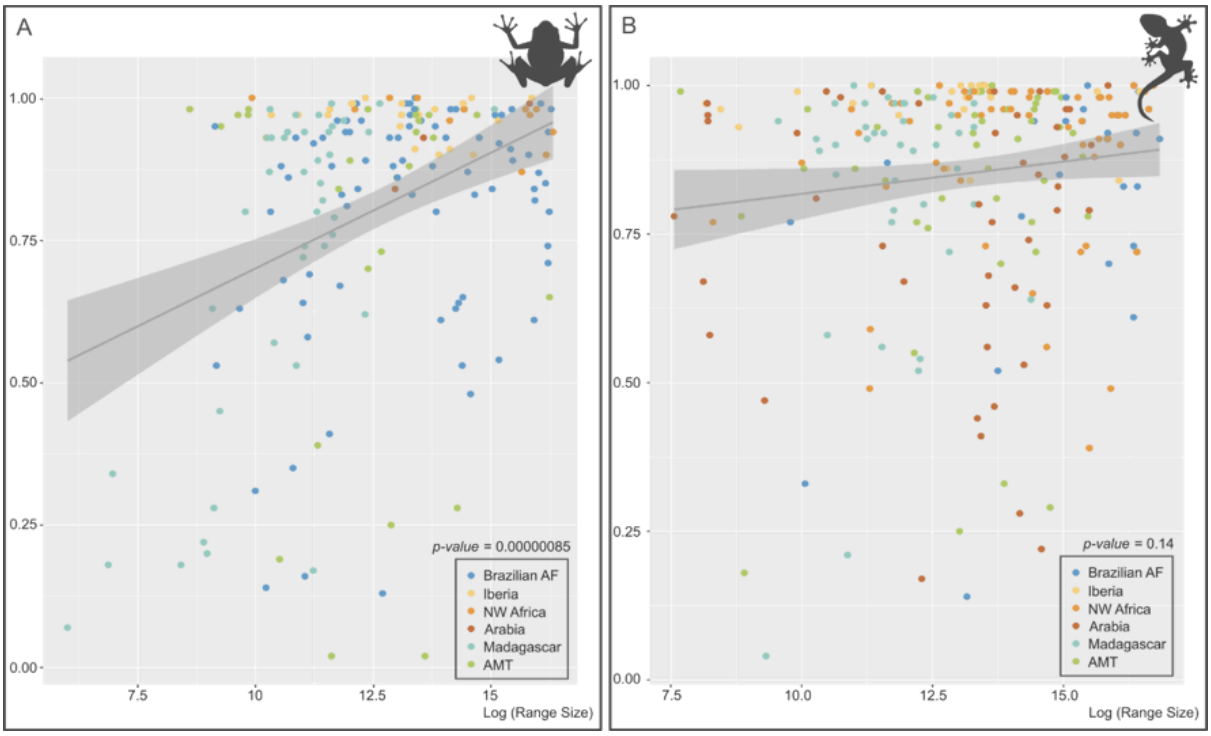


**Figure S5.** Relationship between range representativeness and log-transformed species range size in amphibians (A) and reptiles (B). Colours represent regions of occurrence.

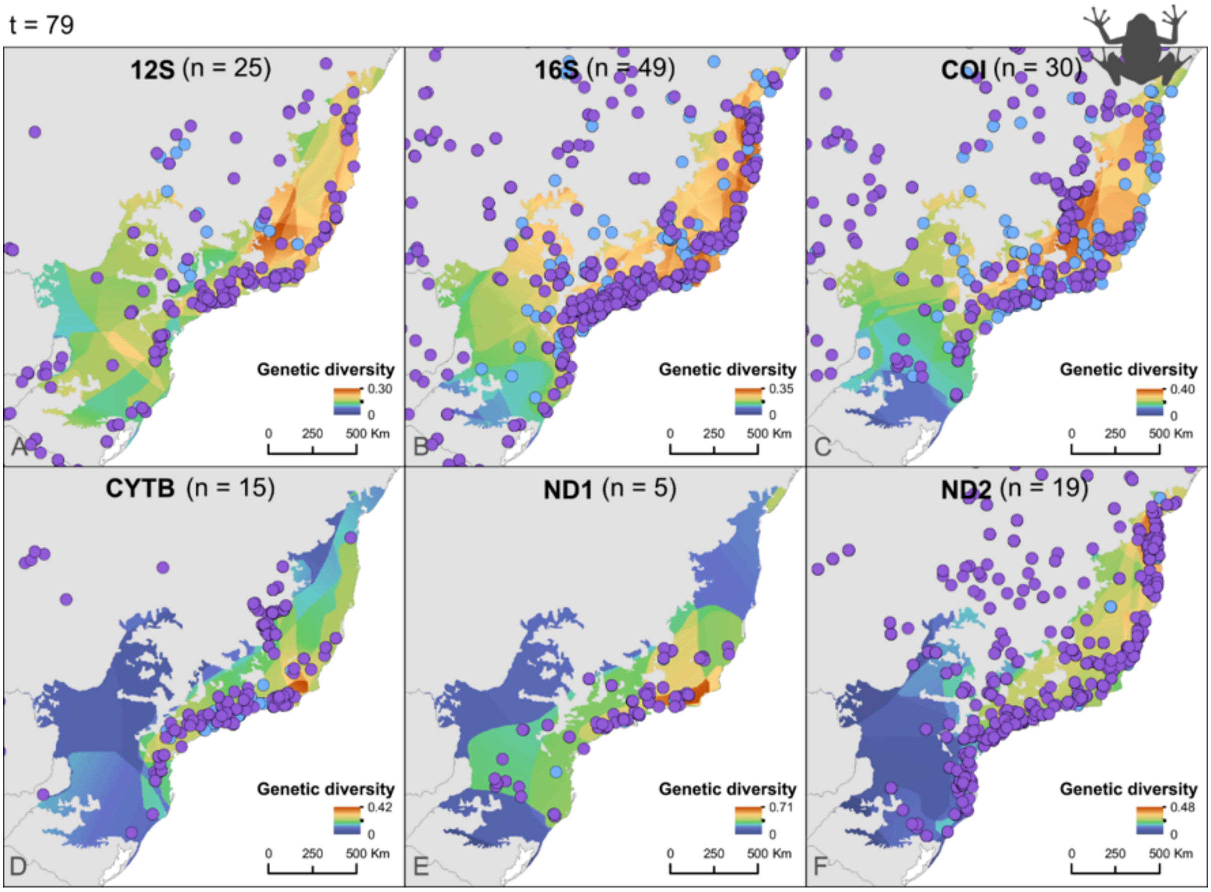


**Figure S6.** Mean nucleotide diversity for (A) 12S, (B) 16S, (C) COI, (D) CYTB, (E) ND1, and (F) ND2 genetic markers, and spatial distribution of mtDNA amphibian samples across the Brazilian AF. (n) is the number of species represented by the plotted marker, (t) is the total number of species retained in the final dataset for this region, as indicated above the figure. Blue dots represent geographic data retrieved through automated approaches (NCBI-provided coordinates or locality descriptors). Purple dots represent geographic data that required manual extraction from the literature (coordinates or locality descriptors reported in publications).

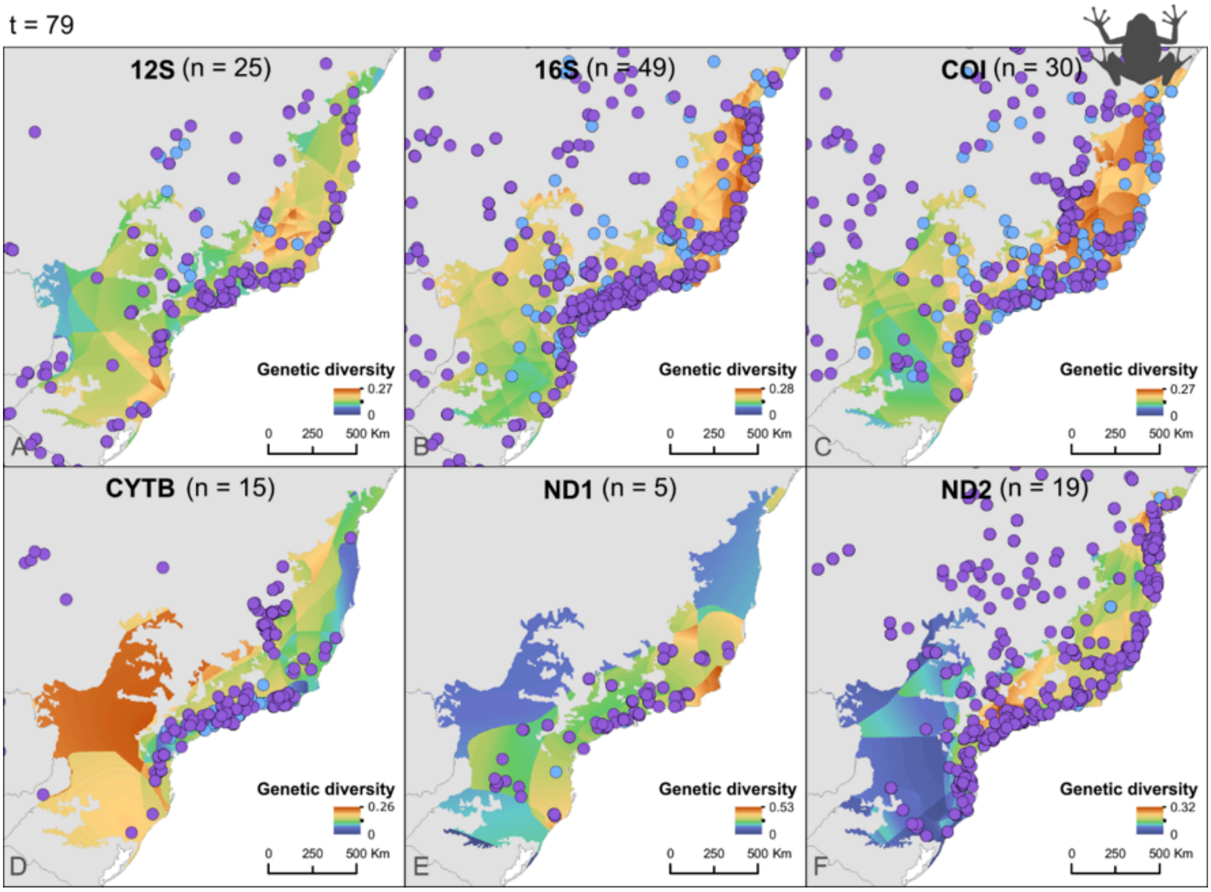


**Figure S7.** Standard error of mean nucleotide diversity for (A) 12S, (B) 16S, (C) COI, (D) CYTB, (E) ND1, and (F) ND2 genetic markers, and spatial distribution of mtDNA amphibian samples across the Brazilian AF. (n) is the number of species represented by the plotted marker, (t) is the total number of species retained in the final dataset for this region, as indicated above the figure. Blue dots represent geographic data retrieved through automated approaches (NCBI-provided coordinates or locality descriptors). Purple dots represent geographic data that required manual extraction from the literature (coordinates or locality descriptors reported in publications).

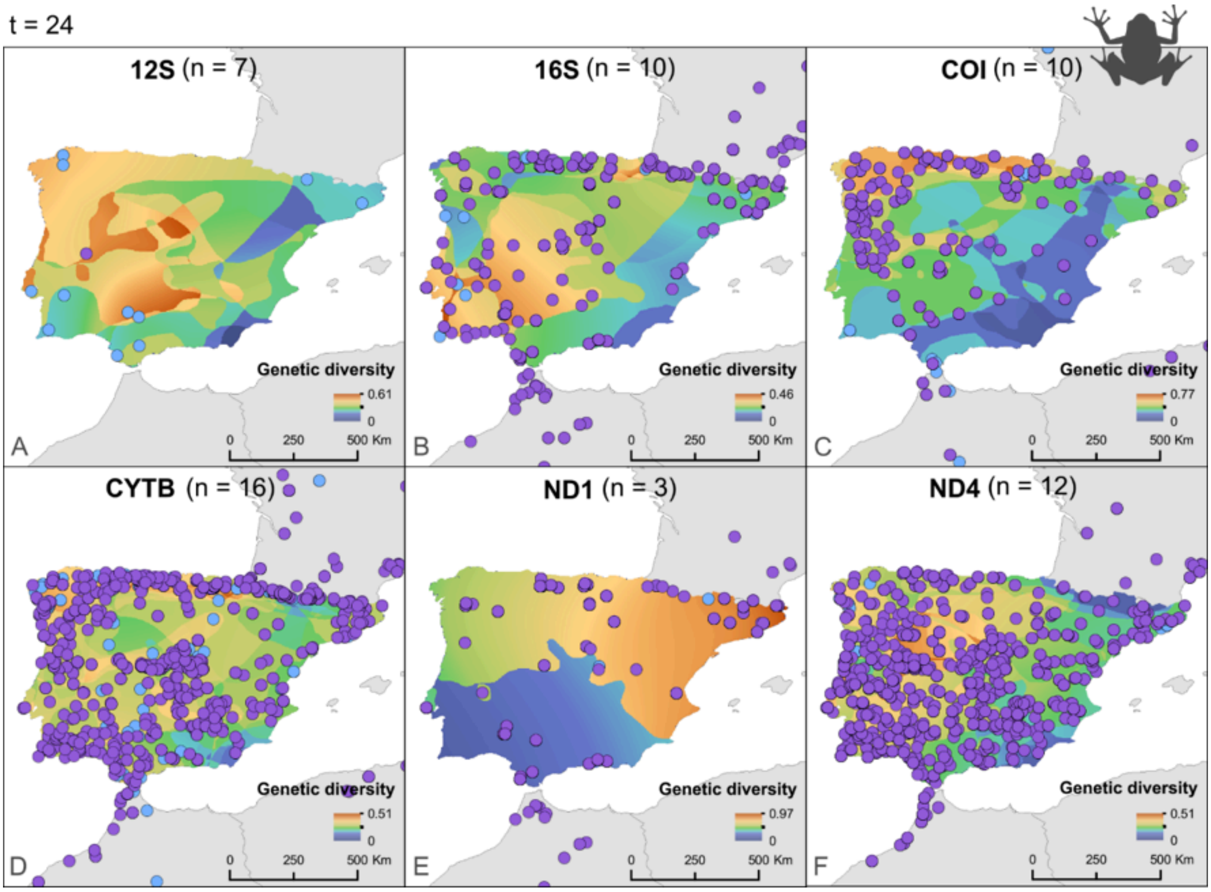


**Figure S8.** Mean nucleotide diversity for (A) 12S, (B) 16S, (C) COI, (D) CYTB, (E) ND1, and (F) ND4 genetic markers, and spatial distribution of mtDNA amphibian samples across the Iberian Peninsula. (n) is the number of species represented by the plotted marker, (t) is the total number of species retained in the final dataset for this region, as indicated above the figure. Blue dots represent geographic data retrieved through automated approaches (NCBI-provided coordinates or locality descriptors). Purple dots represent geographic data that required manual extraction from the literature (coordinates or locality descriptors reported in publications).

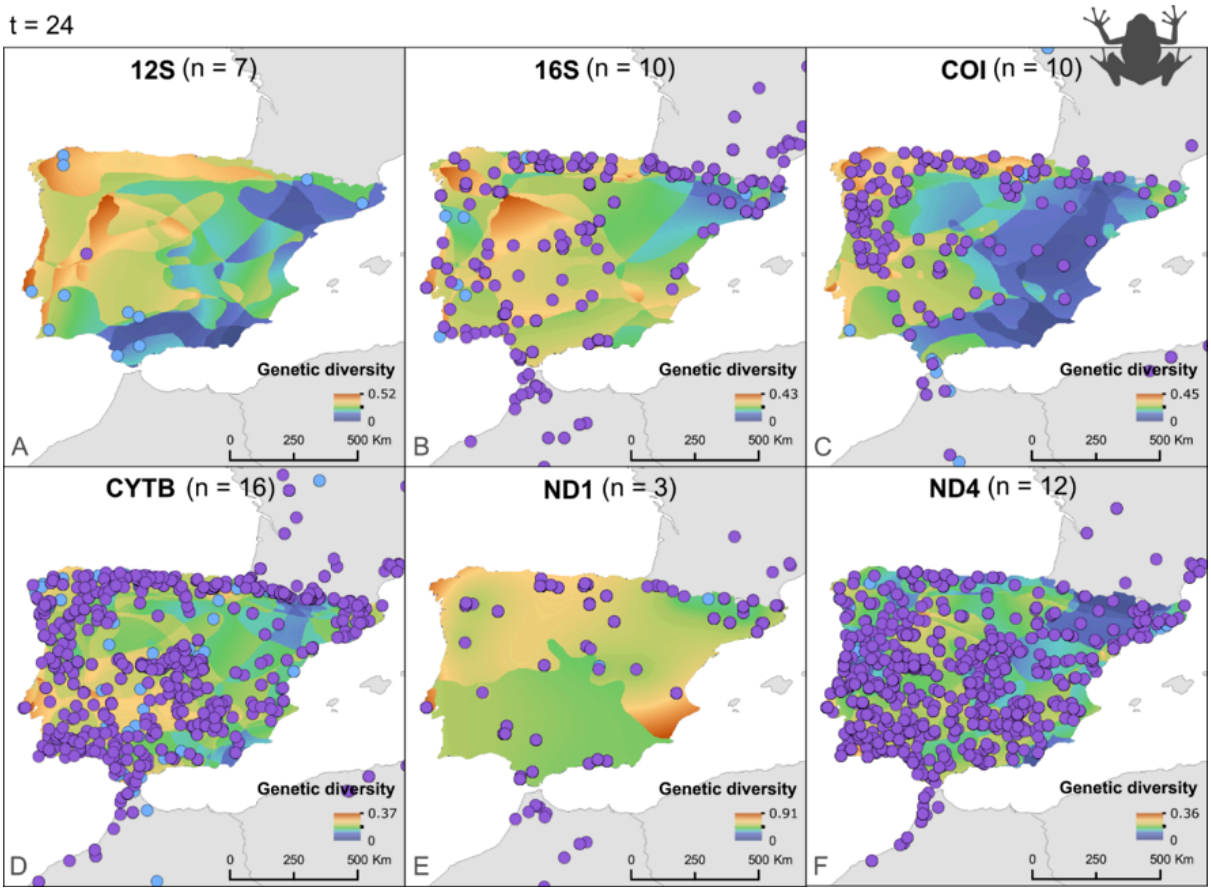


**Figure S9.** Standard error of mean nucleotide diversity for (A) 12S, (B) 16S, (C) COI, (D) CYTB, (E) ND1, and (F) ND4 genetic markers, and spatial distribution of mtDNA amphibian samples across the Iberian Peninsula. (n) is the number of species represented by the plotted marker, (t) is the total number of species retained in the final dataset for this region, as indicated above the figure. Blue dots represent geographic data retrieved through automated approaches (NCBI-provided coordinates or locality descriptors). Purple dots represent geographic data that required manual extraction from the literature (coordinates or locality descriptors reported in publications).

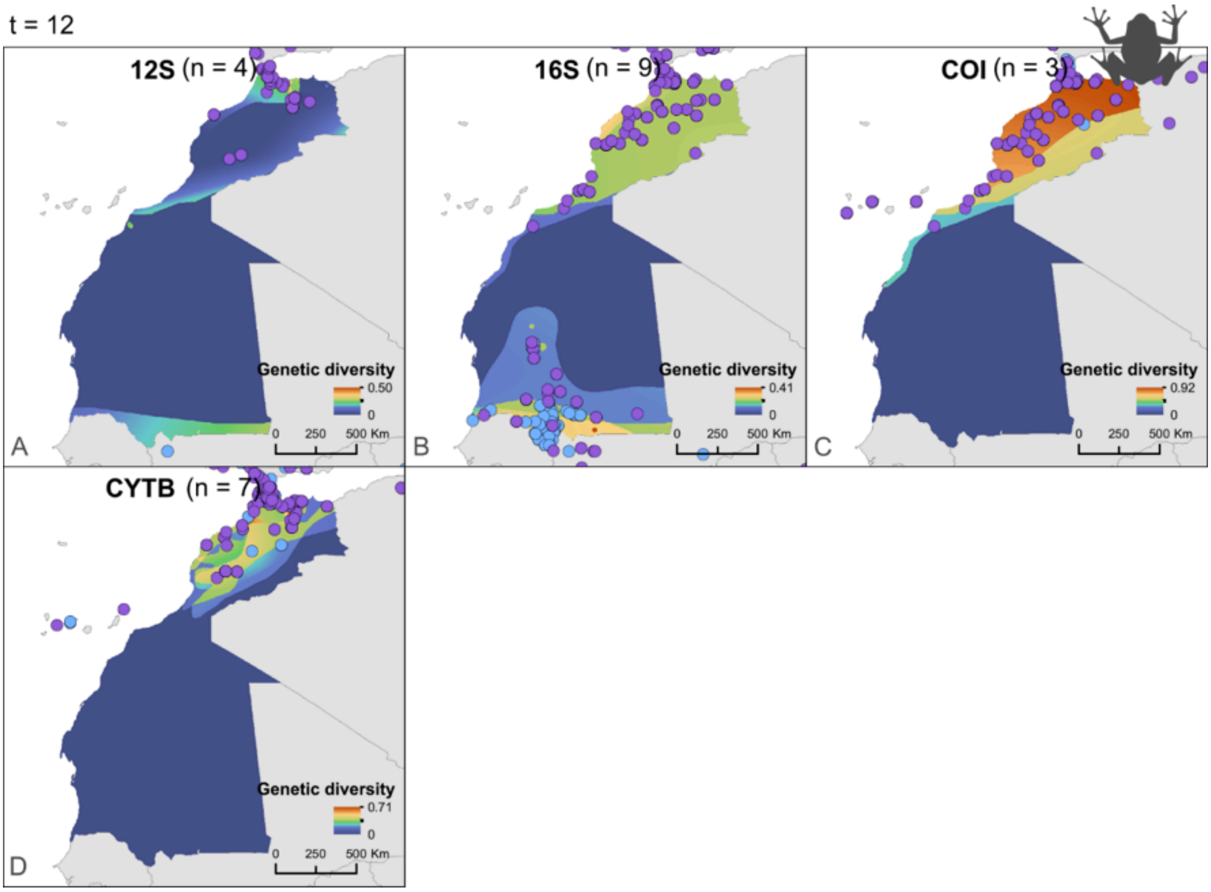


**Figure S10.** Mean nucleotide diversity for (A) 12S, (B) 16S, (C) COI, and (D) CYTB genetic markers, and spatial distribution of mtDNA amphibian samples across NW Africa. (n) is the number of species represented by the plotted marker, (t) is the total number of species retained in the final dataset for this region, as indicated above the figure. Blue dots represent geographic data retrieved through automated approaches (NCBI-provided coordinates or locality descriptors). Purple dots represent geographic data that required manual extraction from the literature (coordinates or locality descriptors reported in publications).

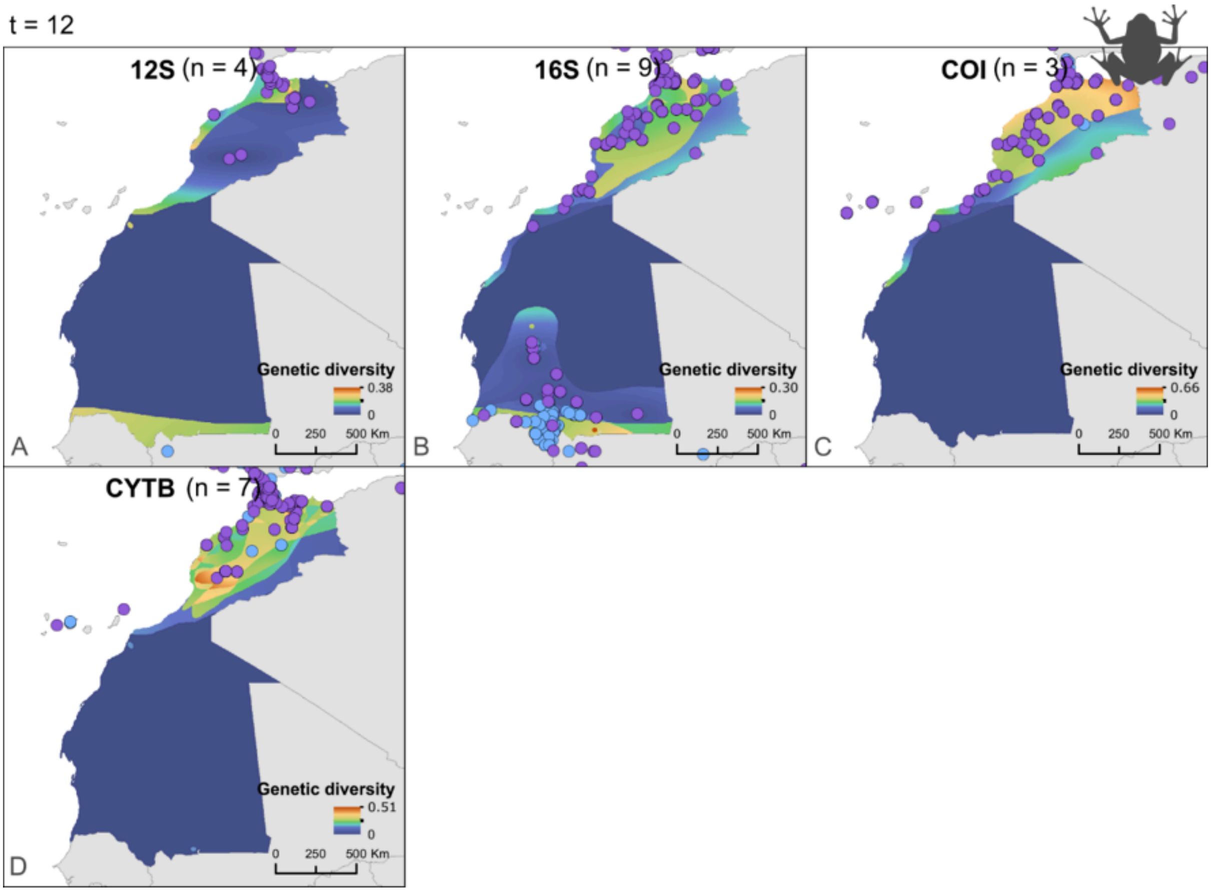


**Figure S11.** Standard error of mean nucleotide diversity for (A) 12S, (B) 16S, (C) COI, and (D) CYTB genetic markers, and spatial distribution of mtDNA amphibian samples across NW Africa. (n) is the number of species represented by the plotted marker, (t) is the total number of species retained in the final dataset for this region, as indicated above the figure. Blue dots represent geographic data retrieved through automated approaches (NCBI-provided coordinates or locality descriptors). Purple dots represent geographic data that required manual extraction from the literature (coordinates or locality descriptors reported in publications).

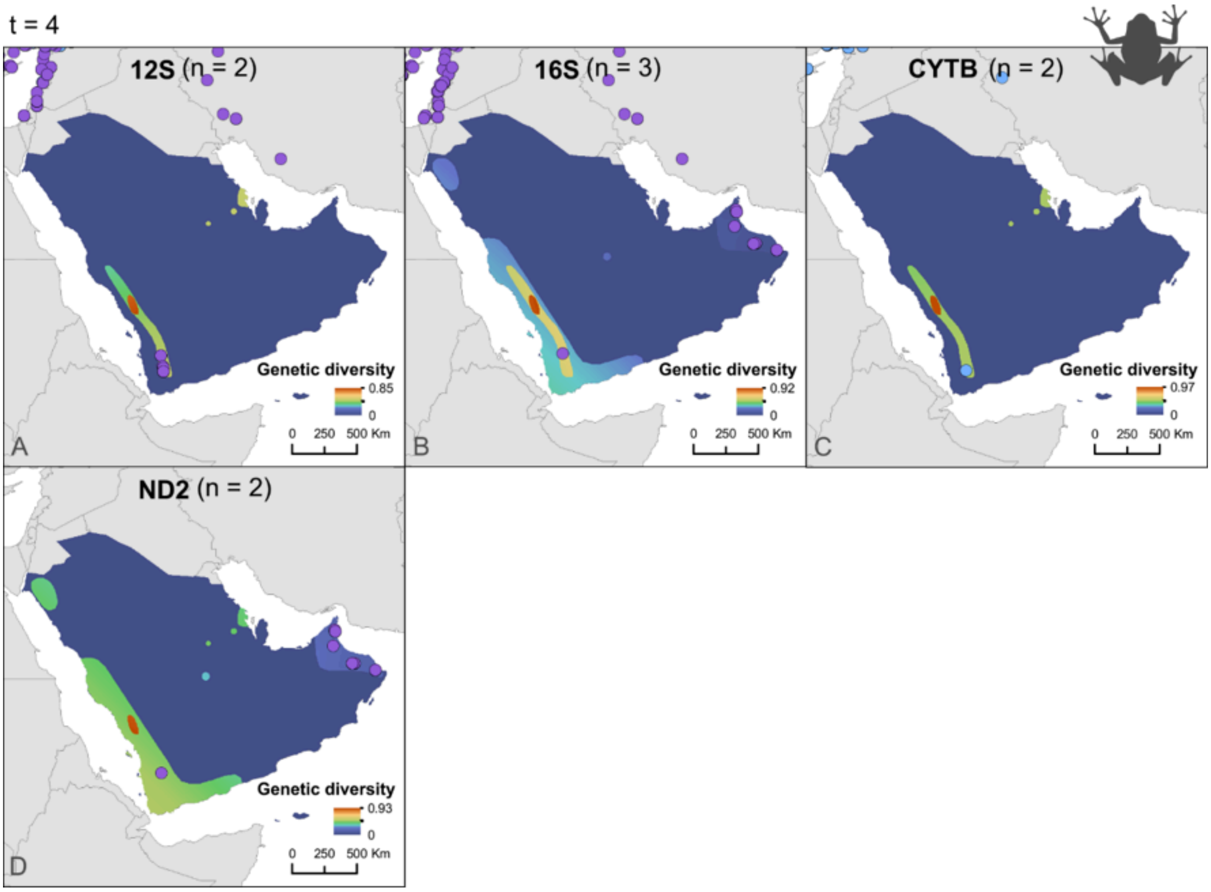


**Figure S12.** Mean nucleotide diversity for (A) 12S, (B) 16S, (C) CYTB, and (D) ND2 genetic markers, and spatial distribution of mtDNA amphibian samples across the Arabian Peninsula. (n) is the number of species represented by the plotted marker, (t) is the total number of species retained in the final dataset for this region, as indicated above the figure. Blue dots represent geographic data retrieved through automated approaches (NCBI-provided coordinates or locality descriptors). Purple dots represent geographic data that required manual extraction from the literature (coordinates or locality descriptors reported in publications).

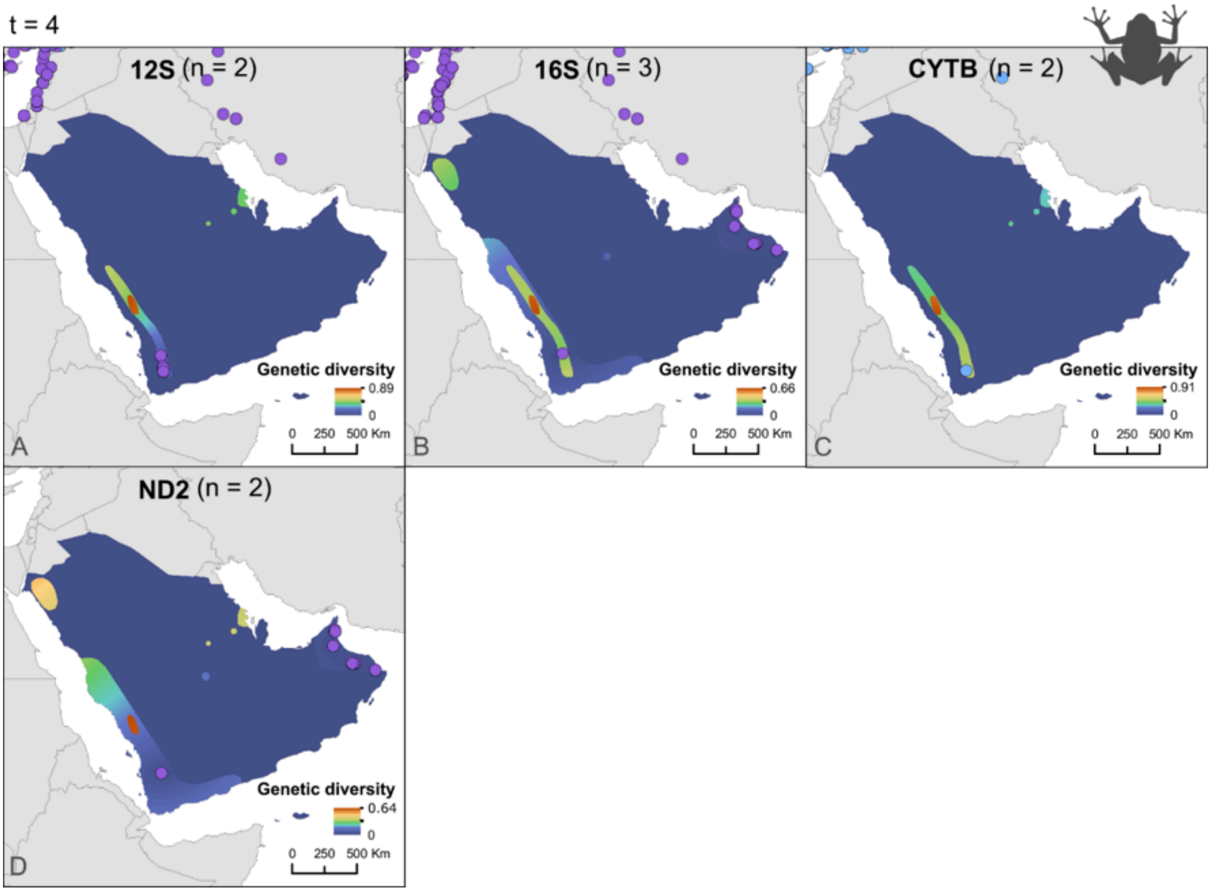


**Figure S13.** Standard error of mean nucleotide diversity for (A) 12S, (B) 16S, (C) CYTB, and (D) ND2 genetic markers, and spatial distribution of mtDNA amphibian samples across the Arabian Peninsula. (n) is the number of species represented by the plotted marker, (t) is the total number of species retained in the final dataset for this region, as indicated above the figure. Blue dots represent geographic data retrieved through automated approaches (NCBI-provided coordinates or locality descriptors). Purple dots represent geographic data that required manual extraction from the literature (coordinates or locality descriptors reported in publications).

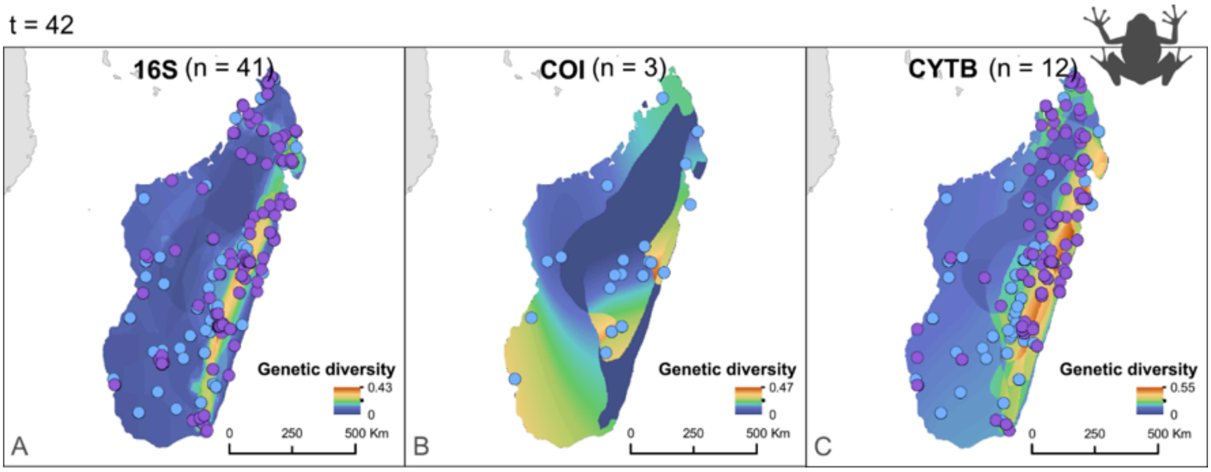


**Figure S14.** Mean nucleotide diversity for (A) 16S, (B) COI, and (C) CYTB genetic markers, and spatial distribution of mtDNA amphibian samples across Madagascar. (n) is the number of species represented by the plotted marker, (t) is the total number of species retained in the final dataset for this region, as indicated above the figure. Blue dots represent geographic data retrieved through automated approaches (NCBI-provided coordinates or locality descriptors). Purple dots represent geographic data that required manual extraction from the literature (coordinates or locality descriptors reported in publications).

t = 42

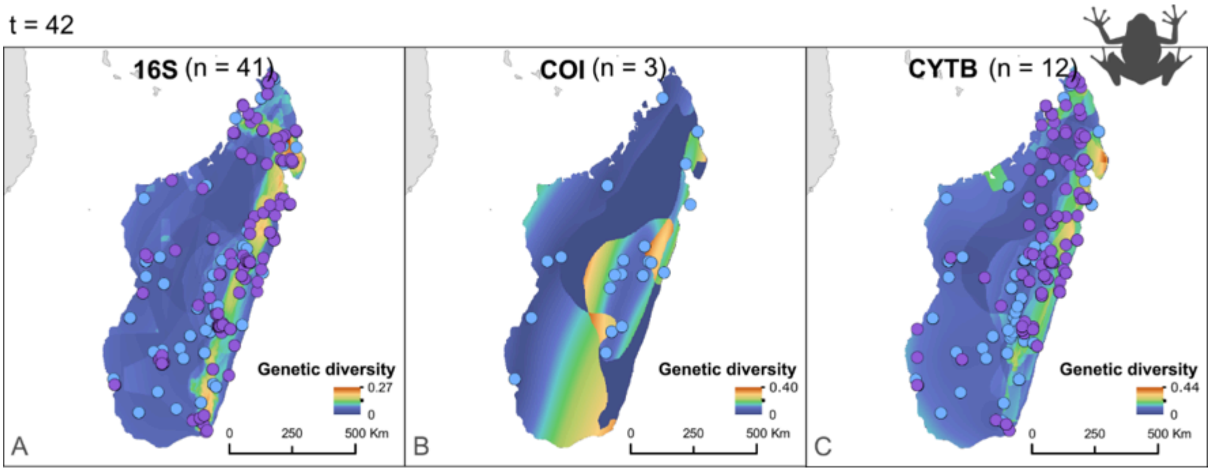


**Figure S15.** Standard error of mean nucleotide diversity for (A) 16S, (B) COI, and (C) CYTB genetic markers, and spatial distribution of mtDNA amphibian samples across Madagascar. (n) is the number of species represented by the plotted marker, (t) is the total number of species retained in the final dataset for this region, as indicated above the figure. Blue dots represent geographic data retrieved through automated approaches (NCBI-provided coordinates or locality descriptors). Purple dots represent geographic data that required manual extraction from the literature (coordinates or locality descriptors reported in publications).

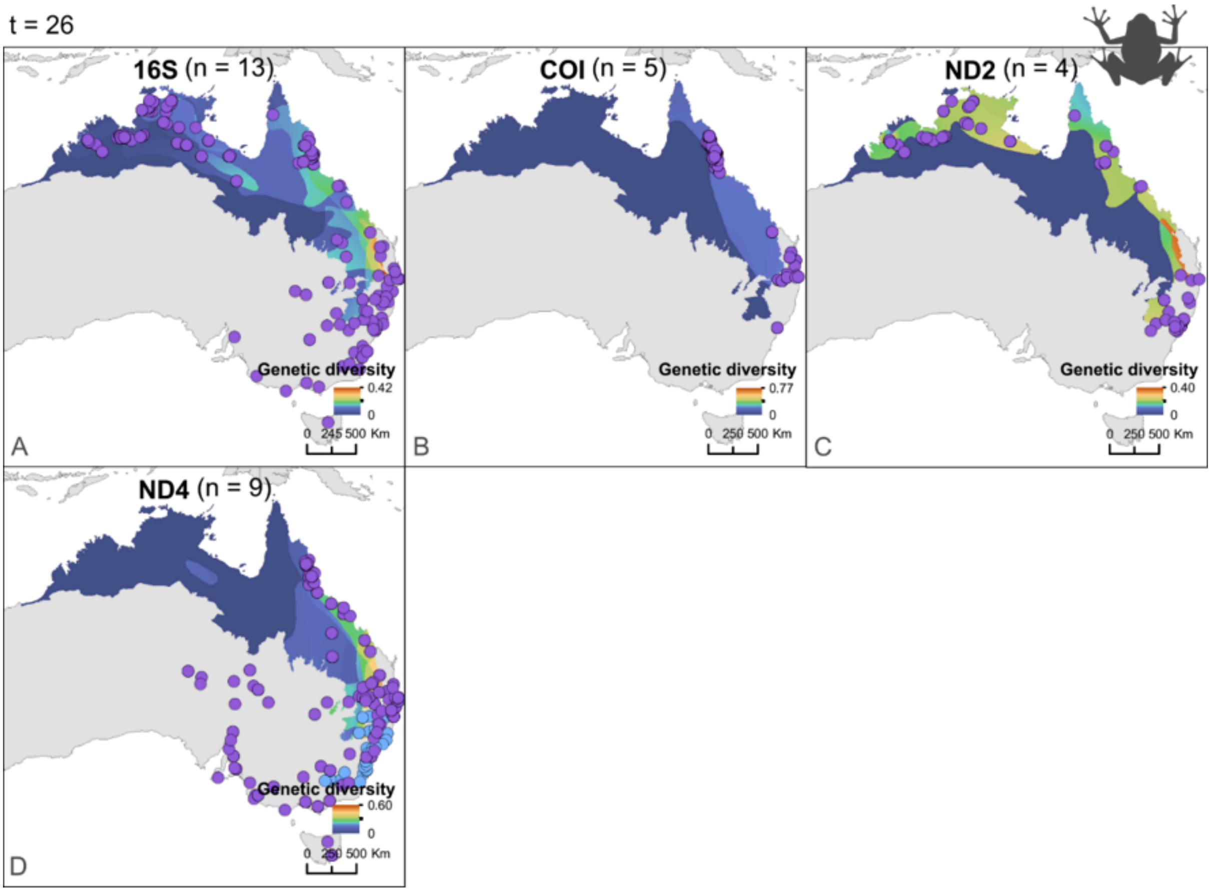


**Figure S16.** Mean nucleotide diversity for (A) 16S, (B) COI, (C) ND2, and (D) ND4 genetic markers, and spatial distribution of mtDNA amphibian samples across AMT. (n) is the number of species represented by the plotted marker, (t) is the total number of species retained in the final dataset for this region, as indicated above the figure. Blue dots represent geographic data retrieved through automated approaches (NCBI-provided coordinates or locality descriptors). Purple dots represent geographic data that required manual extraction from the literature (coordinates or locality descriptors reported in publications).

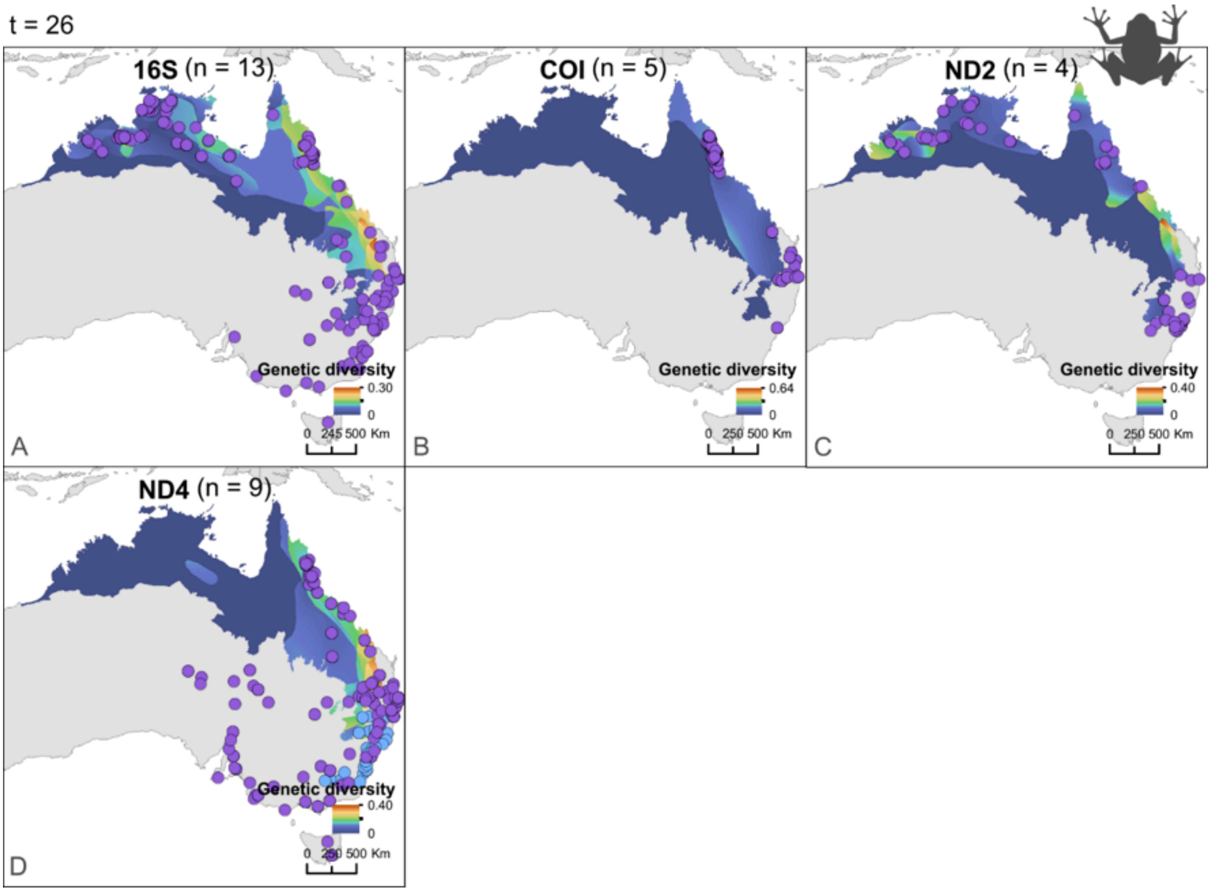


**Figure S17.** Standard error of mean nucleotide diversity for (A) 16S, (B) COI, (C) ND2, and (D) ND4 genetic markers, and spatial distribution of mtDNA amphibian samples across AMT. (n) is the number of species represented by the plotted marker, (t) is the total number of species retained in the final dataset for this region, as indicated above the figure. Blue dots represent geographic data retrieved through automated approaches (NCBI-provided coordinates or locality descriptors). Purple dots represent geographic data that required manual extraction from the literature (coordinates or locality descriptors reported in publications).

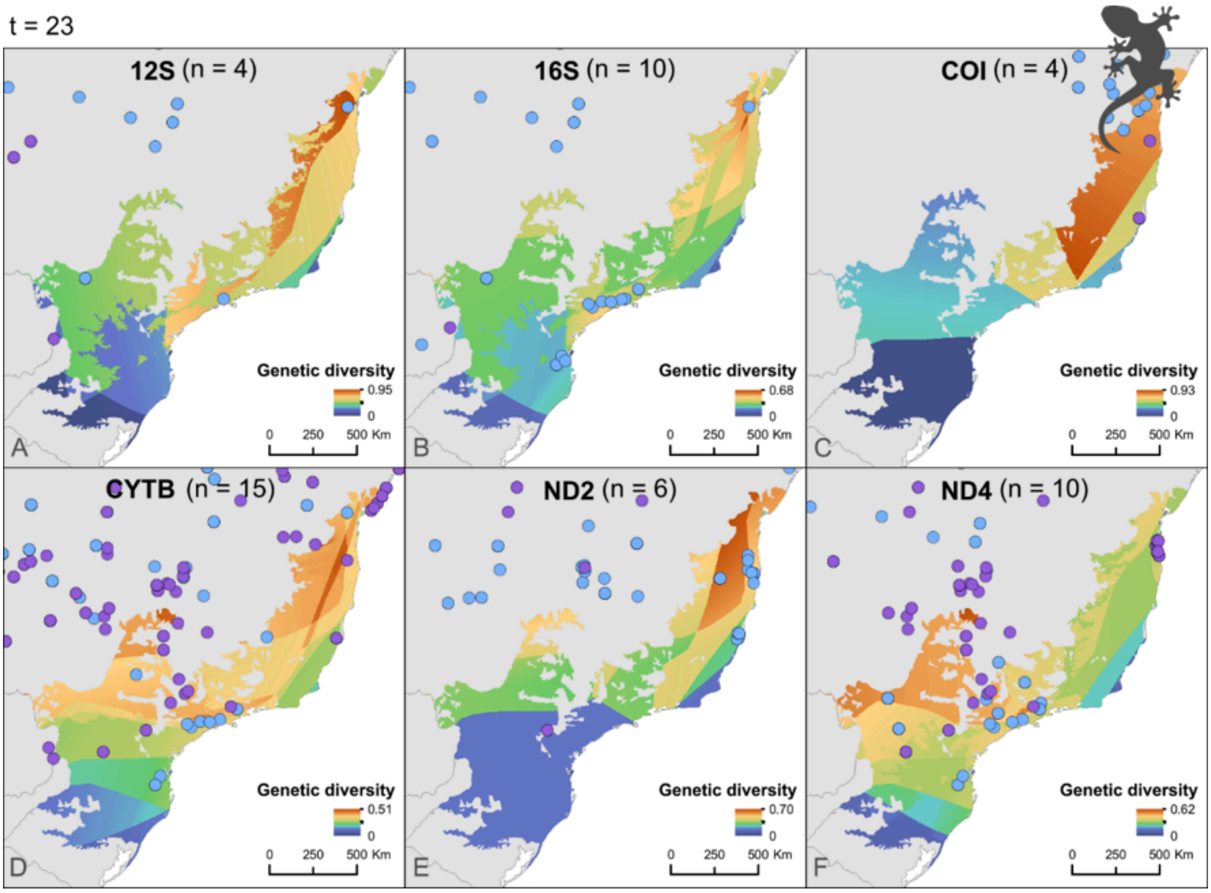


**Figure S18.** Mean nucleotide diversity for (A) 12S, (B) 16S, (C) COI, (D) CYTB, (E) ND2 and (F) ND4 genetic markers, and spatial distribution of mtDNA reptile samples across the Brazilian AF. (n) is the number of species represented by the plotted marker, (t) is the total number of species retained in the final dataset for this region, as indicated above the figure. Blue dots represent geographic data retrieved through automated approaches (NCBI-provided coordinates or locality descriptors). Purple dots represent geographic data that required manual extraction from the literature (coordinates or locality descriptors reported in publications).

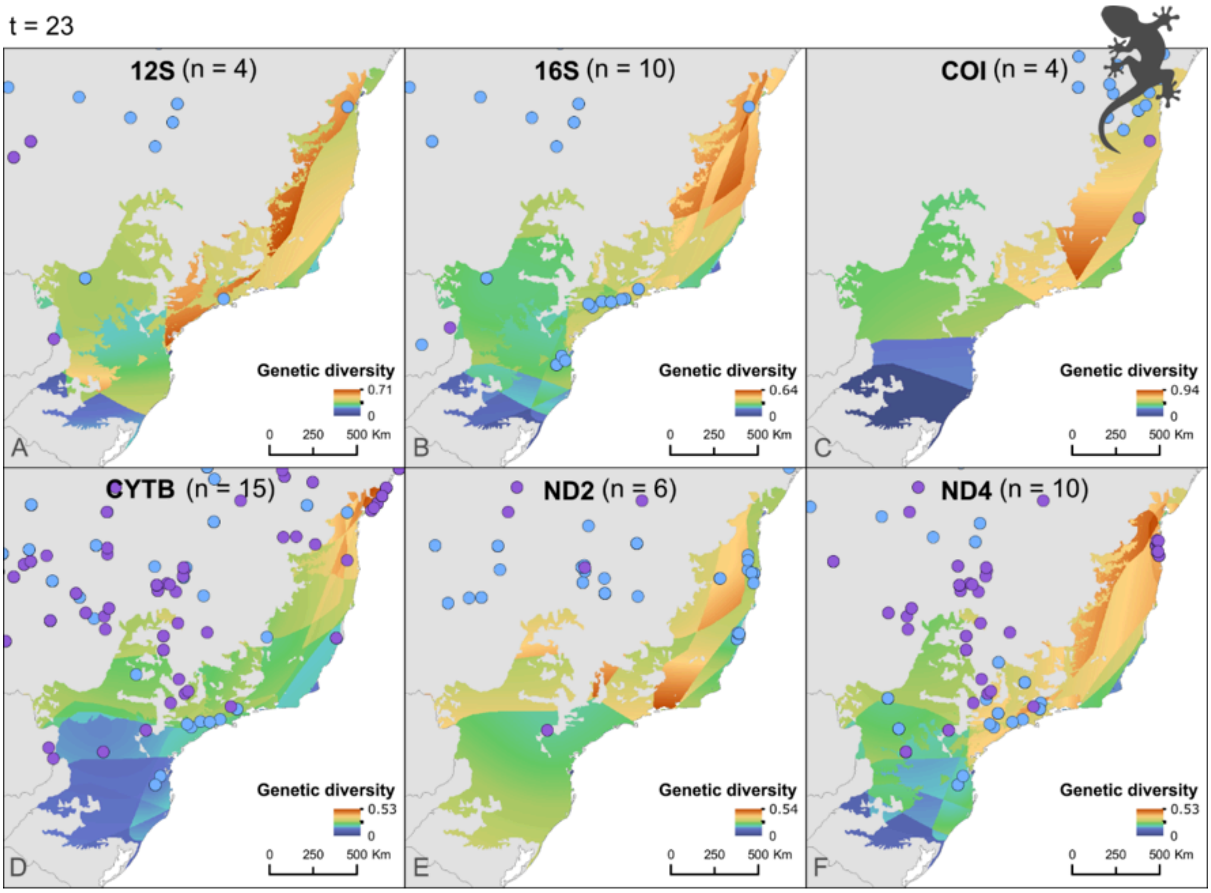


**Figure S19.** Standard error of mean nucleotide diversity for (A) 12S, (B) 16S, (C) COI, (D) CYTB, (E) ND2 and (F) ND4 genetic markers, and spatial distribution of mtDNA reptile samples across the Brazilian AF. (n) is the number of species represented by the plotted marker, (t) is the total number of species retained in the final dataset for this region, as indicated above the figure. Blue dots represent geographic data retrieved through automated approaches (NCBI-provided coordinates or locality descriptors). Purple dots represent geographic data that required manual extraction from the literature (coordinates or locality descriptors reported in publications).

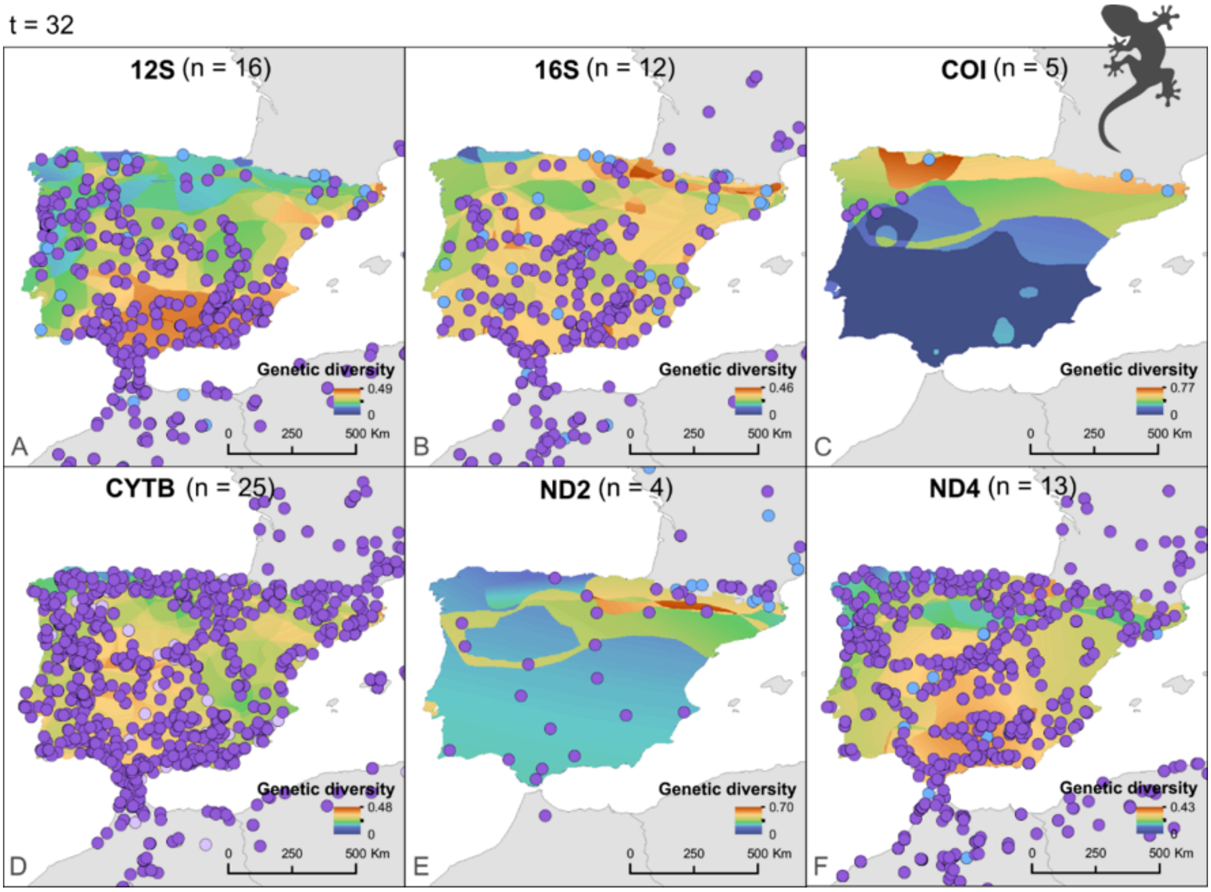


**Figure S20.** Mean nucleotide diversity for (A) 12S, (B) 16S, (C) COI, (D) CYTB, (E) ND2, and (F) ND4 genetic markers, and spatial distribution of mtDNA reptile samples across the Iberian Peninsula. (n) is the number of species represented by the plotted marker, (t) is the total number of species retained in the final dataset for this region, as indicated above the figure. Blue dots represent geographic data retrieved through automated approaches (NCBI-provided coordinates or locality descriptors). Purple dots represent geographic data that required manual extraction from the literature (coordinates or locality descriptors reported in publications).

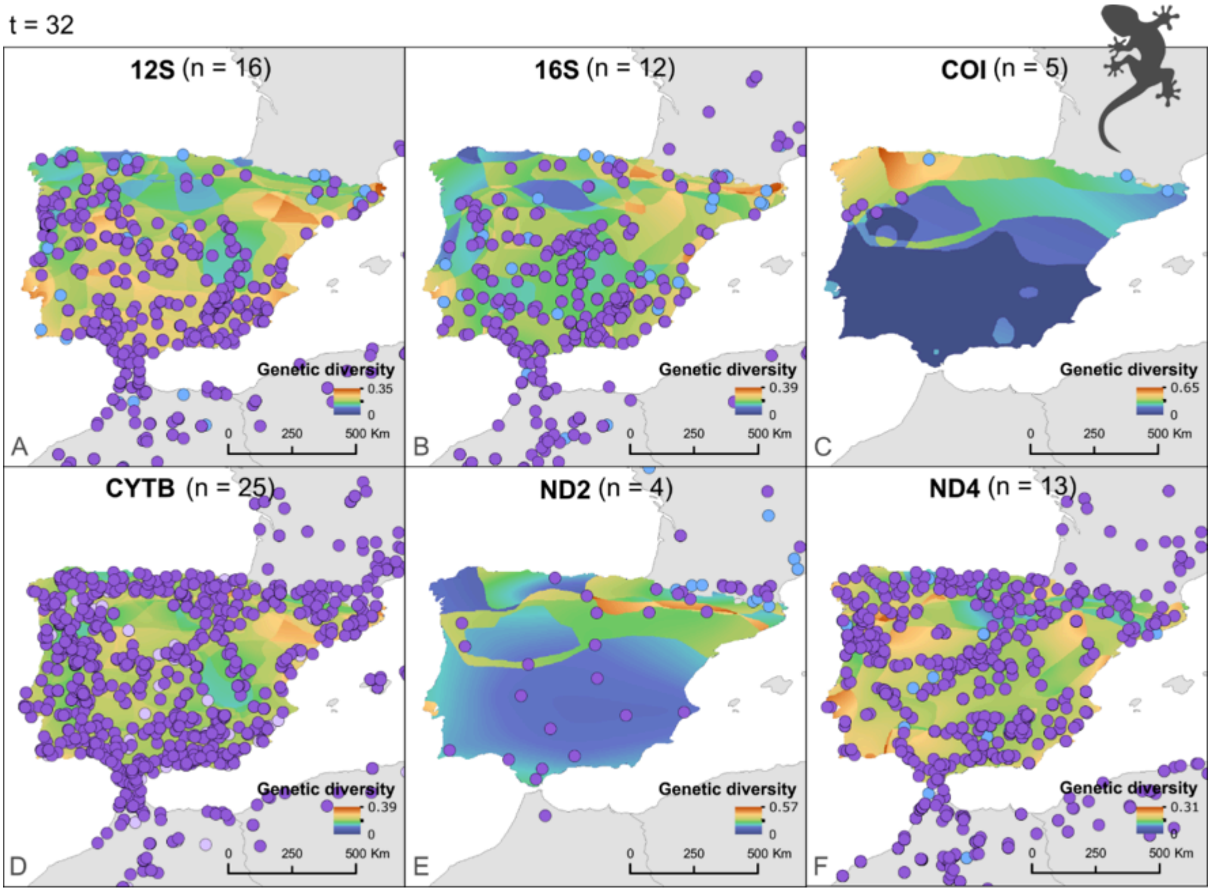


**Figure S21.** Standard error of mean nucleotide diversity for (A) 12S, (B) 16S, (C) COI, (D) CYTB, (E) ND2, and (F) ND4 genetic markers, and spatial distribution of mtDNA reptile samples across the Iberian Peninsula. (n) is the number of species represented by the plotted marker, (t) is the total number of species retained in the final dataset for this region, as indicated above the figure. Blue dots represent geographic data retrieved through automated approaches (NCBI-provided coordinates or locality descriptors). Purple dots represent geographic data that required manual extraction from the literature (coordinates or locality descriptors reported in publications).

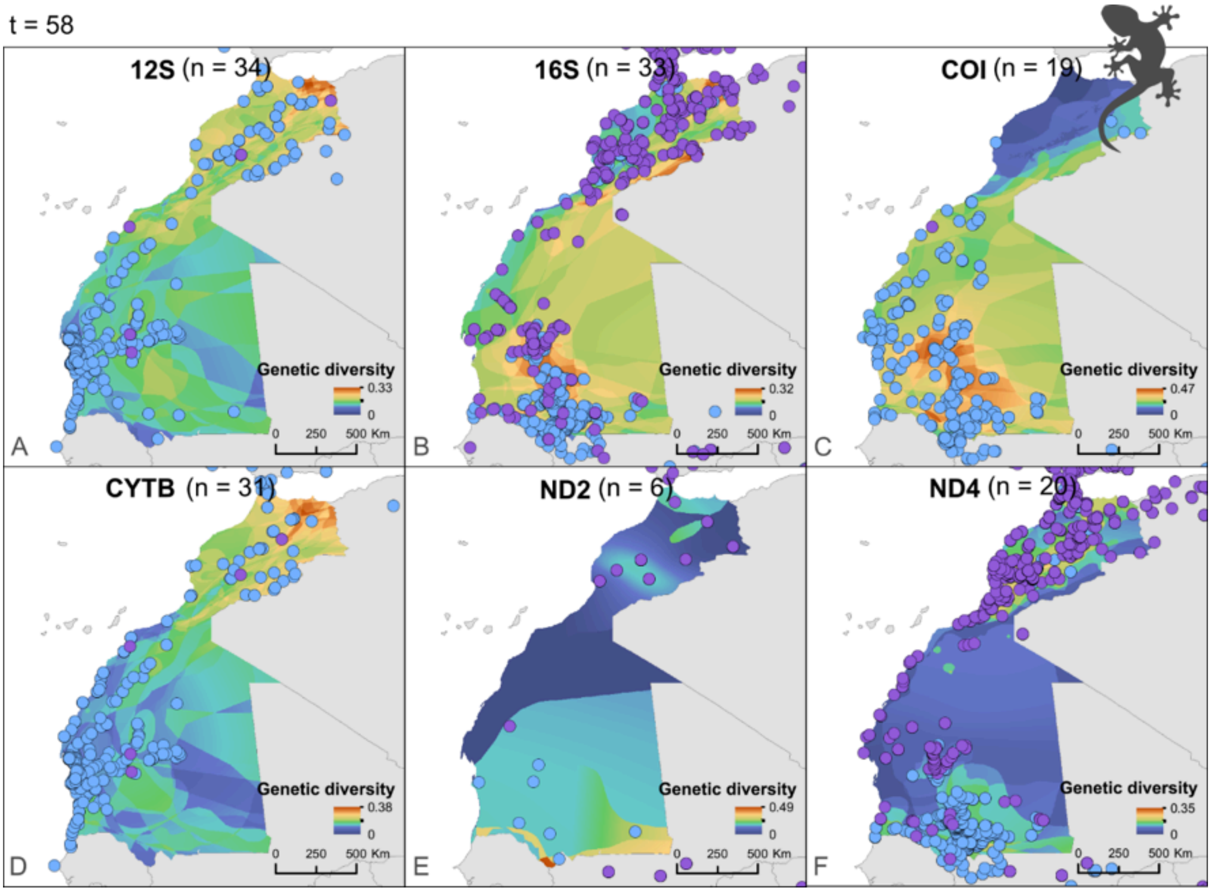


**Figure S22.** Mean nucleotide diversity for (A) 12S, (B) 16S, (C) COI, (D) CYTB, (E) ND2, and (F) ND4 genetic markers, and spatial distribution of mtDNA reptile samples across NW Africa. (n) is the number of species represented by the plotted marker, (t) is the total number of species retained in the final dataset for this region, as indicated above the figure. Blue dots represent geographic data retrieved through automated approaches (NCBI-provided coordinates or locality descriptors). Purple dots represent geographic data that required manual extraction from the literature (coordinates or locality descriptors reported in publications).

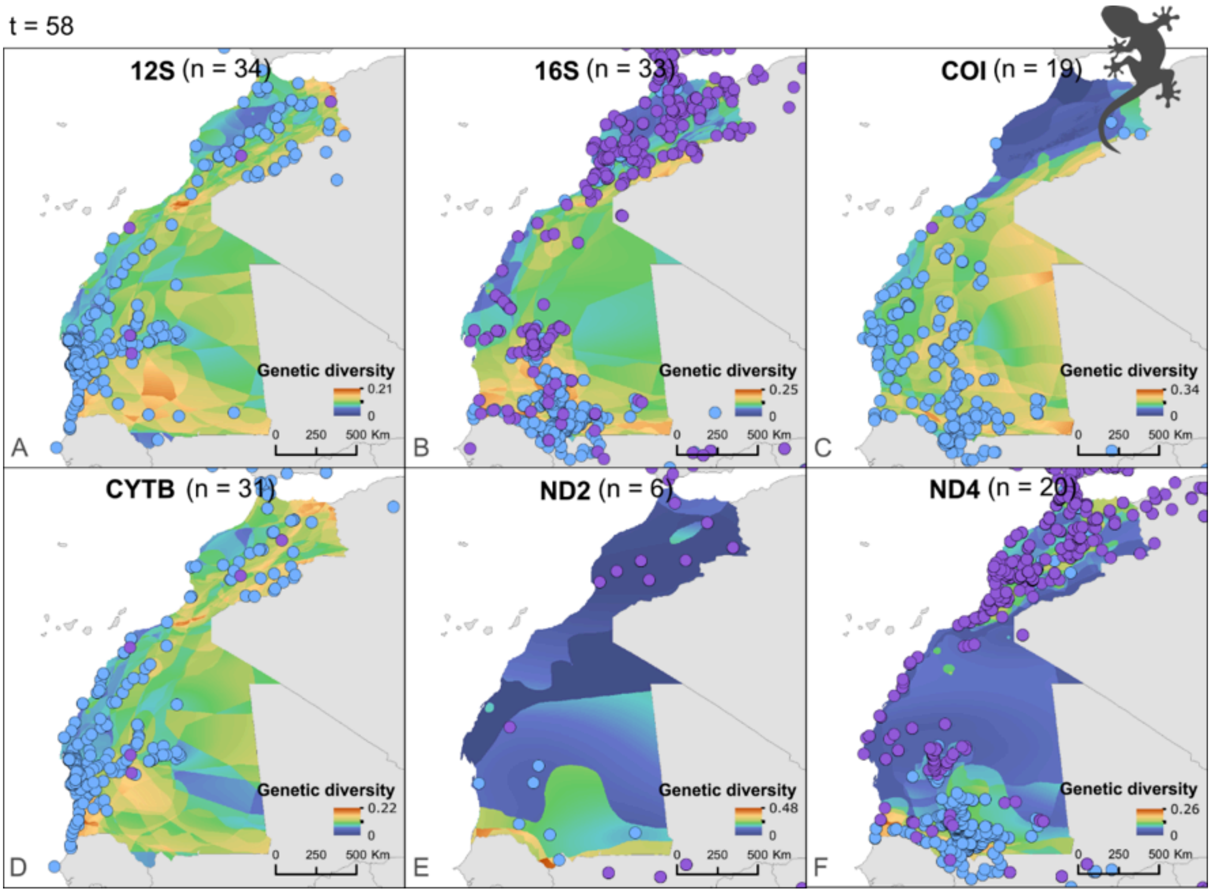


**Figure S23.** Standard error of mean nucleotide diversity for (A) 12S, (B) 16S, (C) COI, (D) CYTB, (E) ND2, and (F) ND4 genetic markers, and spatial distribution of mtDNA reptile samples across NW Africa. (n) is the number of species represented by the plotted marker, (t) is the total number of species retained in the final dataset for this region, as indicated above the figure. Blue dots represent geographic data retrieved through automated approaches (NCBI-provided coordinates or locality descriptors). Purple dots represent geographic data that required manual extraction from the literature (coordinates or locality descriptors reported in publications).

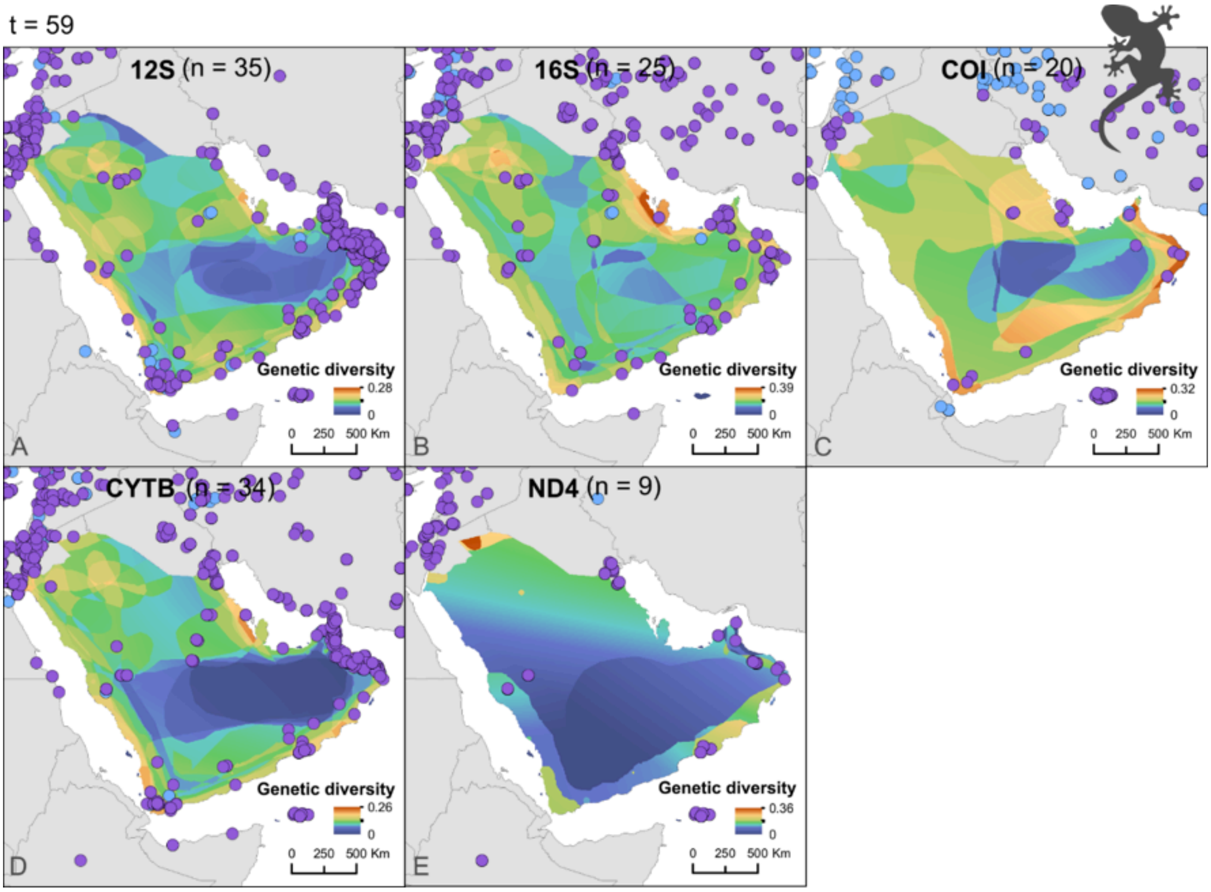


**Figure S24.** Mean nucleotide diversity for (A) 12S, (B) 16S, (C) COI, (D) CYTB, and (E) ND4 genetic markers, and spatial distribution of mtDNA reptile samples across the Arabian Peninsula. (n) is the number of species represented by the plotted marker, (t) is the total number of species retained in the final dataset for this region, as indicated above the figure. Blue dots represent geographic data retrieved through automated approaches (NCBI-provided coordinates or locality descriptors). Purple dots represent geographic data that required manual extraction from the literature (coordinates or locality descriptors reported in publications).

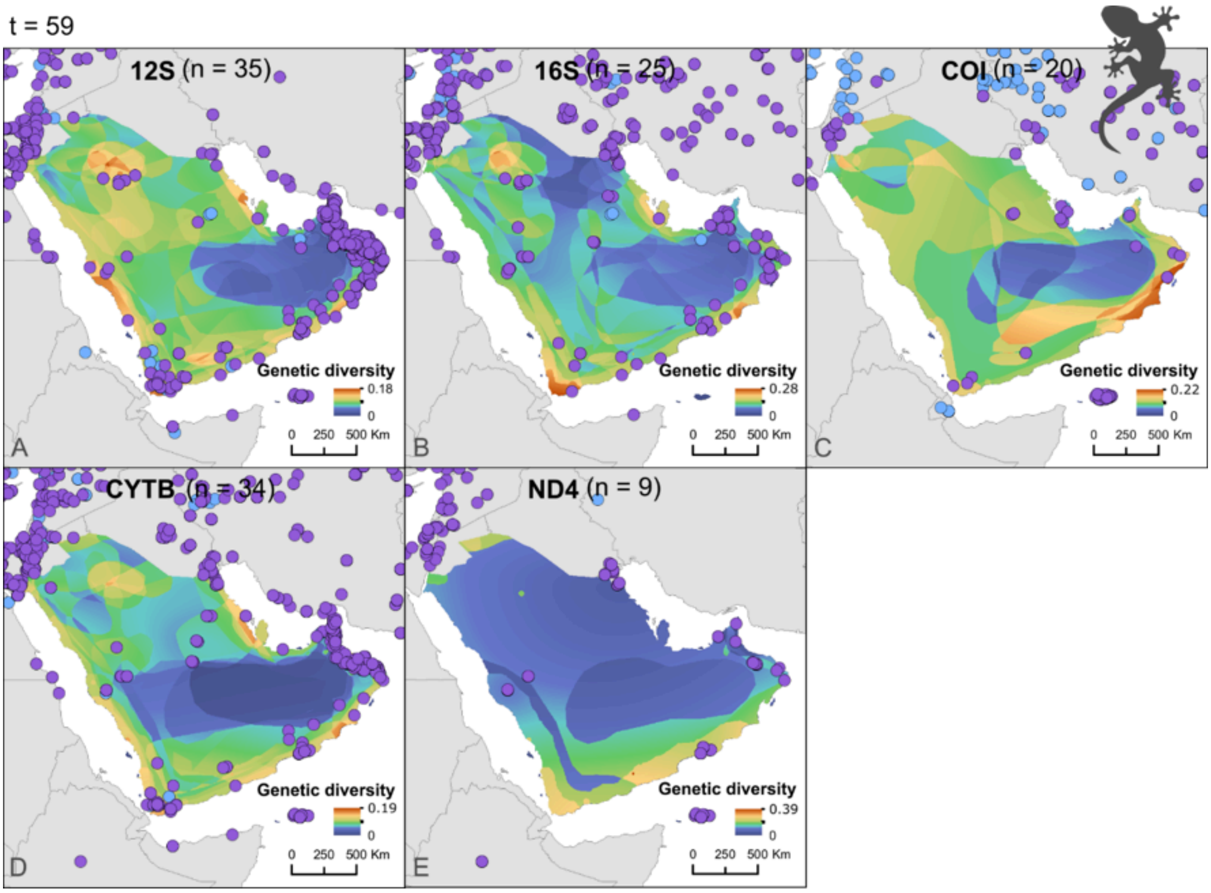


**Figure S25.** Standard error of mean nucleotide diversity for (A) 12S, (B) 16S, (C) COI, (D) CYTB, and (E) ND4 genetic markers, and spatial distribution of mtDNA reptile samples across the Arabian Peninsula. (n) is the number of species represented by the plotted marker, (t) is the total number of species retained in the final dataset for this region, as indicated above the figure. Blue dots represent geographic data retrieved through automated approaches (NCBI-provided coordinates or locality descriptors). Purple dots represent geographic data that required manual extraction from the literature (coordinates or locality descriptors reported in publications).

t = 43

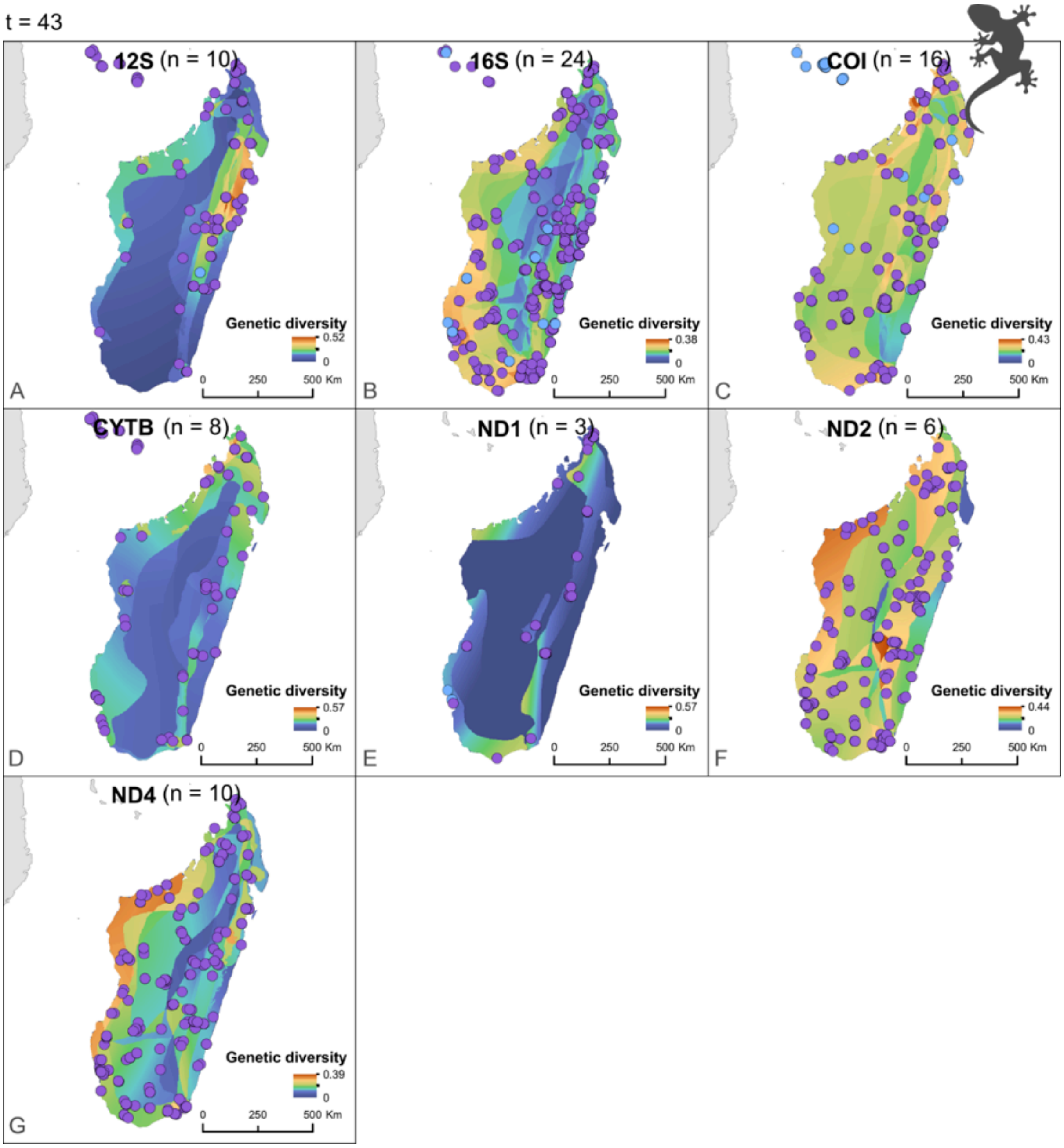


**Figure S26.** Mean nucleotide diversity for (A) 12S, (B) 16S, (C) COI, (D) CYTB, (E) ND1, (F) ND2 and (G) ND4 genetic markers, and spatial distribution of mtDNA reptile samples across Madagascar. (n) is the number of species represented by the plotted marker, (t) is the total number of species retained in the final dataset for this region, as indicated above the figure. Blue dots represent geographic data retrieved through automated approaches (NCBI-provided coordinates or locality descriptors). Purple dots represent geographic data that required manual extraction from the literature (coordinates or locality descriptors reported in publications).

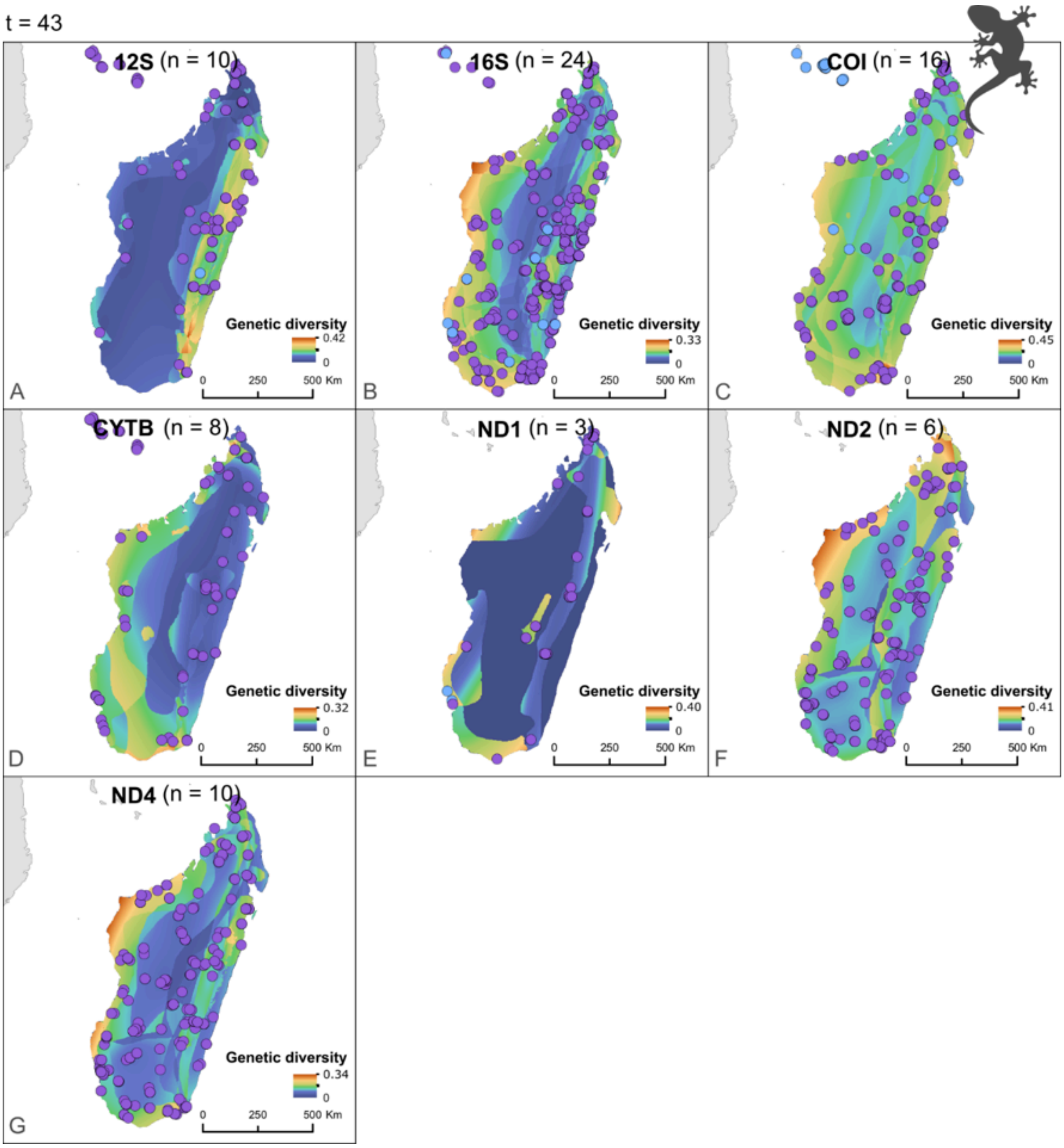


**Figure S27.** Standard error of mean nucleotide diversity for (A) 12S, (B) 16S, (C) COI, (D) CYTB, (E) ND1, (F) ND2 and (G) ND4 genetic markers, and spatial distribution of mtDNA reptile samples across Madagascar. (n) is the number of species represented by the plotted marker, (t) is the total number of species retained in the final dataset for this region, as indicated above the figure. Blue dots represent geographic data retrieved through automated approaches (NCBI-provided coordinates or locality descriptors). Purple dots represent geographic data that required manual extraction from the literature (coordinates or locality descriptors reported in publications).

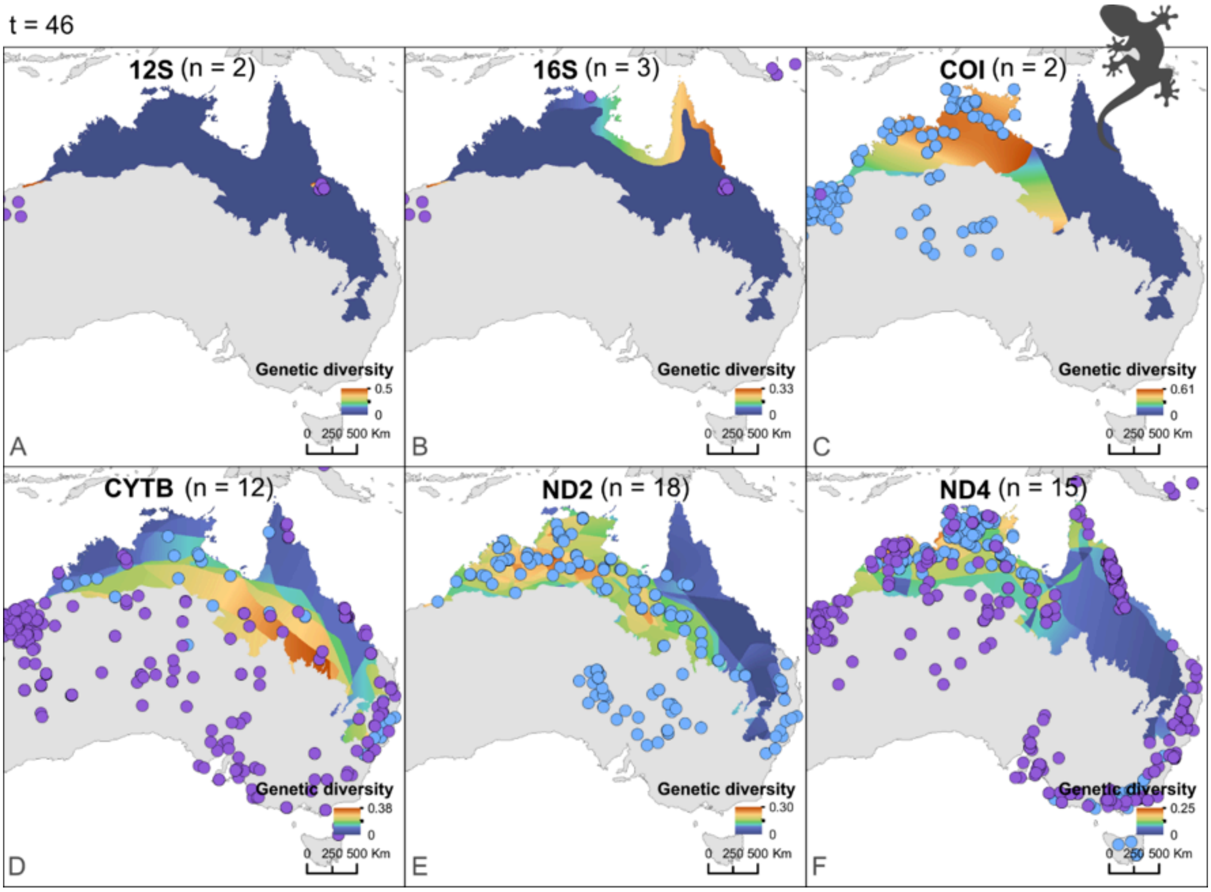


**Figure S28.** Mean nucleotide diversity for (A) 12S, (B) 16S, (C) COI, (D) CYTB, (E) ND2 and (F) ND4 genetic markers, and spatial distribution of mtDNA reptile samples across AMT. (n) is the number of species represented by the plotted marker, (t) is the total number of species retained in the final dataset for this region, as indicated above the figure. Blue dots represent geographic data retrieved through automated approaches (NCBI-provided coordinates or locality descriptors). Purple dots represent geographic data that required manual extraction from the literature (coordinates or locality descriptors reported in publications).

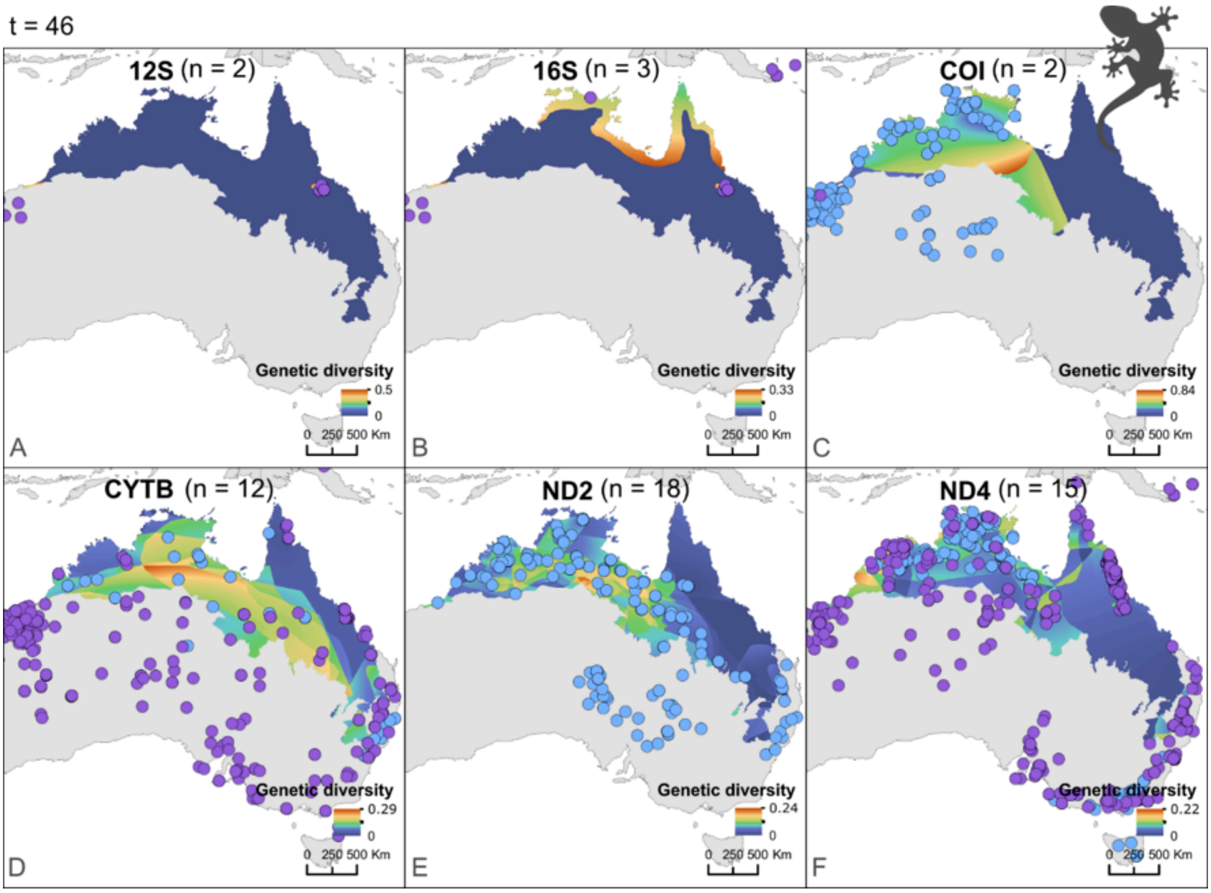


**Figure S29.** Standard error of mean nucleotide diversity for (A) 12S, (B) 16S, (C) COI, (D) CYTB, (E) ND2 and (F) ND4 genetic markers, and spatial distribution of mtDNA reptile samples across AMT. (n) is the number of species represented by the plotted marker, (t) is the total number of species retained in the final dataset for this region, as indicated above the figure. Blue dots represent geographic data retrieved through automated approaches (NCBI-provided coordinates or locality descriptors). Purple dots represent geographic data that required manual extraction from the literature (coordinates or locality descriptors reported in publications).
